# Stepwise developmental mimicry generates proximal-biased kidney organoids

**DOI:** 10.1101/2024.06.28.601028

**Authors:** Jack Schnell, Zhen Miao, MaryAnne Achieng, Connor C. Fausto, Victoria Wang, Faith De Kuyper, Matthew E. Thornton, Brendan Grubbs, Junhyong Kim, Nils O. Lindström

## Abstract

The kidney maintains body fluid homeostasis by reabsorbing essential compounds and excreting waste. Proximal tubule cells, crucial for renal reabsorption of a range of sugars, ions, and amino acids, are highly susceptible to damage, leading to pathologies necessitating dialysis and kidney transplants. While human pluripotent stem cell-derived kidney organoids are used for modeling renal development, disease, and injury, the formation of proximal nephron cells in these 3D structures is incomplete. Here, we describe how to drive the development of proximal tubule precursors in kidney organoids by following a blueprint of *in vivo* human nephrogenesis. Transient manipulation of the PI3K signaling pathway activates Notch signaling in the early nephron and drives nephrons toward a proximal precursor state. These “proximal-biased” (PB) organoid nephrons proceed to generate proximal nephron precursor cells. Single-cell transcriptional analyses across the organoid nephron differentiation, comparing control and PB types, confirm the requirement of transient Notch signaling for proximal development. Indicative of functional maturity, PB organoids demonstrate dextran and albumin uptake, akin to *in vivo* proximal tubules. Moreover, PB organoids are highly sensitive to nephrotoxic agents, display an injury response, and drive expression of *HAVCR1*/*KIM1*, an early proximal-specific marker of kidney injury. Injured PB organoids show evidence of collapsed tubules, DNA damage, and upregulate the injury-response marker *SOX9*. The PB organoid model therefore has functional relevance and potential for modeling mechanisms underpinning nephron injury. These advances improve the use of iPSC-derived kidney organoids as tools to understand developmental nephrology, model disease, test novel therapeutics, and for understanding human renal physiology.

## INTRODUCTION

Proximal nephron cells are the most abundant cells in the human kidney, are responsible for reabsorbing 65% of the nephron filtrate^1^, and their pathologies are the primary reason patients require dialysis and kidney transplants^2,3^. In spite of their clinical importance, proximal tubule disease mechanisms are poorly understood, and human cell models are required to scrutinize disease origins and etiology. Stem cell-derived human kidney models recapitulating proximal tubule functions would therefore provide a critical tool for studying renal disease and develop new therapeutic approaches.

Directed differentiation protocols coaxing induced pluripotent stem cells (iPSCs) to intermediate mesoderm lineages have led to the development of human kidney-like organoids^4,5^. These models partially replicate developing kidney cell profiles, but do not form mature proximal tubule cells, and the proximal precursor-like cells that do develop exhibit low expression of genes that normally impart nephron-specific physiologies^6–11^. While current organoid models have demonstrated upregulation of specific injury markers such as KIM1/HAVCR1 and γH2AX in LTL^+^ cells^12–14^, the lack of homogenous proximal tubule like cells in organoid limits their utility in studying acute proximal tubular injury and performing proximal nephron-specific drug screens.

Studies performed in mice and organoids show that proximal tubule development is dependent on expression of transcription factor *Hnf4a*/*HNF4A*, whose protein product is required for the normal expression of roughly 300 proximal tubule solute carriers, protein, ion, and substrate transporters, and other proximal nephron functional genes^15–17^. Recently, effort has therefore focused on generating HNF4A^+^ proximal tubule precursors as these would serve as building blocks for generating functional proximal tubules. One strategy has enriched for this cell population by expanding the pool of nephron progenitors in early organoids and subsequently allowed their differentiation^10^, but regardless of the protocol, proximal nephron precursors display relatively low expression of *HNF4A* and HNF4A-dependent genes^6–11,15^.

*In vivo*, proximal precursors emerge in the S-shaped body nephron following a stereotyped and deeply conserved developmental program^18^. In this program, nephron progenitors are gradually recruited from their niche into pretubular aggregates that sequentially undergo epithelial-to-mesenchymal transitions generating epithelial renal vesicles. Complex morphogenetic events form tubular Comma-shaped and thereafter the aforementioned S-shaped body (SSB) nephrons with distal and proximal gene signatures positioned along the emerging distal-to-proximal axial polarity^18–20^. Transcriptionally distinct HNF4A^+^ proximal tubule precursors develop in narrow 2-3 cell-wide populations within each medial domain of SSBs, in a field of Notch ligand JAG1^+^ cells^18,19,21^. Their development is dependent on Notch signaling, as *Notch1* and *Notch2* loss-of-function mice fail to express transcription factor *Hnf1b*, which in turn binds to and is necessary for *Hnf4a* expression^22–24^. In addition to Notch, the development of the proximal-distal nephron axis requires integrated signaling between several pathways as spatial positions in the nascent nephron are known to be driven by Notch, Wnt, BMP, and PI3K signaling, each tuning the formation of various precursor populations^18,25^. As expected, nephrons forming in kidney organoids respond to changes in Wnt signaling^10,26^ in a manner conserved with that shown in mouse kidneys^25^, but there is no clear maturation of cells. This raises the possibility to developmentally program organoid nephrons to form proximal tubule precursors, but such strategies have not been identified and are therefore required.

In this study, we address this challenge by developing a protocol to expand the HNF4A^+^ proximal nephron precursor population within kidney organoids and in individual nephrons. We direct organoid cell differentiation along an *in vivo*-like developmental trajectory proceeding through JAG1^+^/HNF1B^+^ fates and culminating in HNF4A^+^ proximal precursors. Comparative analyses with *in vivo* development show the organoid proximal precursors mature to resemble HNF4A^+^ cells in the capillary loop stage (CLSN) human nephron with emerging physiologies. Proximal-biased (PB) nephrons display expression of solute carriers and transporters and are capable of selectively transporting albumin and dextran. We further leverage the prevalence of proximal structures within the PB organoid model to demonstrate significant upregulation of KIM1/HAVCR1 within HNF4A^+^ organoid nephron tubules in response to nephrotoxic injury. Directly relevant to *in vivo* injury, this response is mosaic and KIM1/HAVCR1^+^, HNF4A^+^ PB organoid nephrons have collapsed apical-basal polarities, show DNA damage, and upregulate injury-response marker *SOX9*. The PB nephron model represents a significant step towards recapitulating kidney development and function in organoid systems and provides a direct strategy to study proximal nephrotoxicity and tubulopathies in a robust human assay.

## RESULTS

### Identifying abnormal kidney organoid developmental programs

To identify differences between developing nephrons in kidneys and organoids that can explain why organoids do not generate maturing proximal cells, we characterized how proximal precursors form *in vivo* in developing human kidneys and compared organoids to this blueprint using single-cell RNA-sequencing and secondary validation for proteins that mark and drive proximal tubule development.

The specification of proximal precursors is consequent to the gradual recruitment of nephron progenitor cells into the forming nephron and signaling pathways that tune differentiation along the progressively emerging proximal-distal axis^21,25^. This process begins with a cellular domain developing in the distal renal vesicle nephron formed by early recruited nephron progenitor cells. The domain is marked by membrane localized JAG1 and nuclear HNF1B and is in direct contact with the ureteric epithelium. It abuts the proximally located WT1^+^ region where cells are actively recruited from the nephron progenitor cell niche (**Figure 1a**). As the renal vesicle develops into a comma-shaped body nephron, the HNF1B^+^/JAG1^+^ domain expands proximally. The initial distal domain downregulates JAG1 to form HNF1B^+^/JAG1^low^ cells, while the proximally expanding domain further upregulates JAG1 to become HNF1B^+^/JAG1^high^, now considered the medial domain of the comma-shaped body. The medial domain also forms a boundary with the WT1^+^ proximal-most domain where the last nephron progenitors are recruited (**Figure 1a; Figure S1a-c**). The nephron is at the comma-shaped nephron stage, subdivided by these markers into three distinguishable domains. As it develops further, HNF1B is additionally upregulated in nuclei positioned at the border between the medial and proximal-most domain, where HNF4A is detected in late comma-shaped/early S-shaped body nephrons (**Figure 1b**). Proximal tubule precursor cells (HNF1B^high^/HNF4A^+^) then develop and downregulate JAG1 as the S-shaped nephron matures into capillary loop stage nephrons, when an elongating tubule forms with a HNF1B^high^/HNF4A^high^/JAG1^-^ proximal cell state (**Figure 1b**). These finds are consistent with single-cell RNA-sequencing of the developing human nephron lineage (**Figure S1a-c**). Proximal precursor cells diverge from other nephron lineages in an early *PAX8^+^* cell population in the pretubular aggregate, which generates *WT1^+^* podocyte precursors, *TFAP2A^+^* distal precursors, and *JAG1^+^* cells that sequentially upregulate *HNF1B* and *HNF4A* (data combined from Tran et al., 2019 *Dev Cell*; Lindstrom et al., 2021 *Dev Cell*)^8,18^ (**Figure S1a-c; Table S1**).

**Figure 1.**
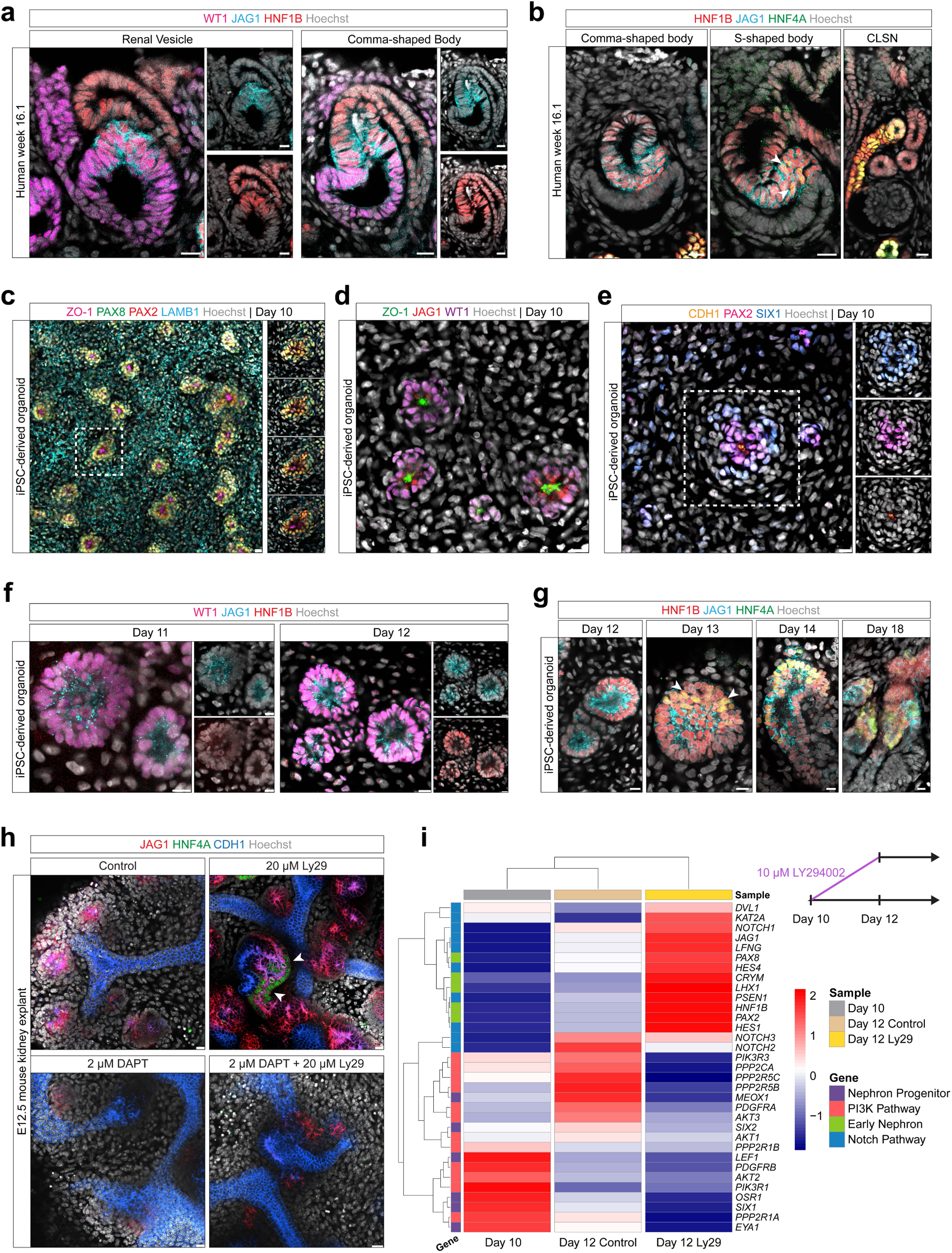
Comparative morphological and transcriptomic analyses of proximal nephron formation *in vivo* and in iPSC-derived kidney organoids. **a-b**. Immunofluorescent antibody stains at defined stages of nephrogenesis. Human kidneys from week 16 of development. (a) represents stages before the onset of HNF4A, while (b) represents the commencement and elongation of the HNF4A^+^ domain. White arrowheads indicate earliest forming HNF4A^+^ cells. Scale bars: 10 microns. **c-e.** Whole-mount immunofluorescent stains of day 10 kidney organoids. Inset in (c) highlights a Z-stack through an individual early forming organoid nephron. Boxed region in (e) shows individual channels. Scale bars: 10 microns. **f-g**. Whole-mount immunofluorescent stains of kidney organoid nephrons. Data in (f) match proteins and stages in (a), while (g) corresponds to (b). White arrowheads indicate earliest forming HNF4A^+^ cells. Scale bars: 10 microns. **h**. *Ex vivo* cultures of dissected E12.5 mouse kidneys in different 48 hr. culture conditions. White arrowheads indicate Hnf4a^+^ cells. Scale bars: 20 microns. **i**. Hierarchical clustering from bulk RNA-sequencing of kidney organoids at differentiation days 10, 12 (control) and 12 (treated for 48 hours with 10 µM LY294002). Genes shown are those that are highlighted in Figure S1g. Legend categorizes genes and timepoints. Timeline of organoid differentiation and those profiled is shown in the top right.

Organoid nephrons form differently. At day 10, kidney organoids consist of individual cell-aggregates positive for nephron progenitor markers (WT1, SIX1) and nephron lineage markers (PAX2, PAX8). These cells coalesce around a forming apical epithelial polarity (CDH1, ZO1) with an accreting basement membrane (LAMB1) (**Figure 1c-e**). As the aggregates further epithelialize (day 11-12), they display uniform deposition of WT1 in nuclei, and JAG1 as puncta and weakly at cell-membranes around the periphery of the forming nephron (**Figure 1f-g**). HNF1B is upregulated between days 11 and 12 and is deposited in nuclei around the periphery of each nephron. At these stages, there are no indications of WT1, JAG1, and HNF1B being distributed along a gradually forming proximal-distal axis as seen *in vivo* (**Figure 1a**), rather the organoid nephrons exhibit a HNF1B^+^/JAG1^+^/WT1^+^ triple-positive cell state (**Figure 1f**). During further differentiation, organoid nephrons generate HNF1B^+^/HNF4A^+^ cells, but these do not transition into a rapidly elongating phase as observed *in vivo* and JAG1 remains strongly detectable (**Figure 1b,g**). Single-cell data from organoids sampled over time and differentiation protocols indicate that the abnormal early and late triple-positive *PAX8^+^/JAG1^+^/WT1^+^* and late *HNF1B^+^/HNF4A^+^/JAG1^+^* cell states are a common trend across models (reanalyzed and combined from several sources)^4–9^ (**Figure S1d-f; Table S2**).

Given the importance of Hnf1b and Hnf4a for normal proximal tubule development, we examined the possibility that organoids abnormally regulate functionally important *Hnf1b/HNF1B* and *Hnf4a/HNF4A*-mediated transcriptional programs and assessed organoid proximal fate development against the *in vivo* developmental blueprint. To identify genes that are co-expressed with each transcription factor *in vivo* (**Figure S1a**), we performed a Pearson expression correlation analysis for each gene and characterized the expression of *HNF1B* and *HNF4A* correlates. The expression of these genes *in vivo* was ordered along a predicted proximal developmental trajectory, and their expression patterns highlight a transition from *HNF1B* correlates in early cells (cluster 9: pretubular aggregate/renal vesicle; e.g., *JAG1*, *HES1*, *KRT8*) through to a gradually more *HNF4A*-correlating signature (clusters 21, 24: S-shaped body; e.g., *CLU*, *DCDC2*, *ANXA4*), and finally genes strongly enriched in the maturing proximal precursors (cluster 20: Capillary loop stage nephron; e.g., *ASS1*, *SLC34A1*, *SLC22A8;* **Figure S2a** (left)). Using this progression as a framework for the temporal *in vivo* sequence of transcriptional events, we compared it to organoid proximal precursor development across models^4–9^ (**Figure S2a** (right) based on **Figure S1d**). *In vitro*, *HNF1B*-correlates such as *JAG1*, *HES1*, *KIF12*, and *KRT8* were detected while other genes, for instance transcription factor *ELF3*, fibroblast growth factor receptor *FGFR4*, serine protease inhibitor *SERPINF2*, and extracellular signaling protein *CYR61* were not. This partial recapitulation of the transcription profile was further reduced for *HNF4A*-correlates, with only 25/46 genes detected across *in vitro* models. Genes coding for a range of protein types were not detected or significantly expressed, for instance solute carriers *SLC22A8*, *SLC5A8*, *SLC16A9*, and transferases and enzymes such as *AGXT2*, *GLYAT*, and *ANPEP*.

To independently examine whether HNF4A-dependent genes are underrepresented in the organoid proximal program, we intersected genes downregulated upon *in vivo Hnf4a* loss-of-function^16^, with transcriptional profiles from detailed expression maps of the adult male and female mouse kidney^27^. We categorized Hnf4a-dependent genes as those expressed only in development and those that persist into the adult functional nephron. 257 genes are significantly downregulated (control vs. mutant kidneys*, p*_adj_ < 0.01, log_2_FC ≥ 1.5)^28^ in postnatal day 0 animals on loss of *Hnf4a* (reanalyzed from Marable et al., 2020 *JASN*)^16^, and 415 genes are enriched in the adult male and female proximal tubules (Kidney Cell Explorer)^27^ (**Figure S2b**). Of the 257 Hnf4a-dependent genes, 124 genes were enriched only in the developing kidney while 133 genes showed persistent expression into functional adult nephrons. To determine whether human orthologs are expressed in the developing human proximal tubule and in organoid models, we intersected our gene lists with single-cell transcriptional data (**Figure S1a,d**). Of the 133 Hnf4a-dependent genes expressed during development and in adult mouse nephrons, 92 human orthologs were detected. 91/92 were also detected in organoids, and 89/92 (96.7%) were detected at greater abundance *in vivo* (**Figure S2c,e; Table S3**). Similarly, of the 124 Hnf4a-dependent genes expressed during development, 81 human orthologs were detected. 80 of these were detected in organoids, but 70/81 (86.4%) were again detected at a higher frequency *in vivo* (Figure S2d,f; Table S4).

These data show that correlates of HNF4A and human orthologs of Hnf4a-dependent genes are expressed infrequently in organoids. This is consistent across organoid models, collectively pointing to abnormal regulation of HNF4A-mediated gene expression.

### Initiating a proximal-forming cell-state in kidney organoids

To develop a strategy to drive proximal precursor development in organoids, we revisited our previous work identifying a relationship between PI3K signaling and Notch ligand Jag1^29^. In mouse kidneys, WNT/β-catenin and PI3K signaling have opposing effects on *Jag1* expression, and pharmacological inhibition of PI3K signaling results in rapid upregulation of *Jag1* in nephron progenitors and early nephrons^25,29^. We therefore tested whether inhibition of PI3K and upregulation *Jag1* could drive a proximal nephron program as this could provide a tool to generate human proximal tubule precursors in kidney organoids. Mouse kidney explants were cultured with PI3K inhibitor LY294002 (Ly29)^30,31^ for 48 hours. This increased Jag1 abundance as expected. Hnf4a was co-detected in Jag1^+^ cells, confirming expression of a *bona fide* proximal precursor marker (**Figure 1h**). To determine if *Hnf4a* expression was dependent on Notch signaling, as suggested by mouse genetic models^32,33^, we treated kidneys with both Ly29 and Notch/gamma-secretase inhibitor DAPT^34,35^. Co-inhibition of PI3K and Notch signaling blocked Hnf4a expression, as did inhibiting Notch on its own (**Figure 1h**). These data confirm an interaction between the PI3K and Notch pathways in generating proximal cell identities in kidneys.

To validate this strategy in kidney organoids, we treated organoids with Ly29 from days 10-12. This is the time when JAG1 is partially upregulated *in vitro* and when nephrons epithelialize. We transcriptionally profiled them before and after treatment using RNA-sequencing. PI3K inhibitor-treated organoids displayed a 1.78-fold increase in *JAG1* expression, and Notch pathway genes *LNFG*, *HES1*, *HES4*, *NOTCH1* were strongly upregulated, as were differentiation markers *PAX8* and *LHX1*. Nephron progenitor markers *SIX1*, *MEOX1*, *OSR1*, *EYA1* were downregulated. Wnt/β-catenin targets *WNT4* and *LGR5* were unchanged (**Figure 1i; Figure S1g; Table S5**). Gene set enrichment analyses confirmed that the Notch pathway was selectively upregulated, PI3K pathway downregulated, and other unrelated pathways, e.g., Hedgehog, were unaffected (**Figure S1h-j**). Within this 48-hr. timeframe, *HNF1B* expression was upregulated 2.88-fold in PI3K inhibited samples, but *HNF4A* was not detected at day 12 (TPM < 1.7; **Figure 1i; Figure S1g; Table S5**). These data show that organoids upregulate *JAG1* and *HNF1B* in response to transient PI3K inhibition.

Upregulation of *JAG1* and *HNF1B* occurred in nephrons alongside elongation (HNF1B 1.64-fold size increase, *p*-value < 0.0001; JAG1, 2.36-fold size increase, *p*-value < 0.0001) and the abundance of HNF1B was also increased within each nephron (1.22-fold increase, *p*-value < 0.0001) compared to controls (**Figure 2a-b**). These changes were dependent on Notch signaling as JAG1^+^/HNF1B^+^ nephrons did not elongate or further upregulate HNF1B when cultured with Notch antagonist DAPT, either with or without Ly29 (**Figure S3a**). The protein encoded by Notch target gene *HES1*, a marker for Notch signaling, was reduced in organoids treated with both Ly29 and DAPT, but was upregulated in organoids treated with only Ly29 (**Figure S3a**). The increase in HES1 on Ly29 treatment persisted through differentiation days 13 and 14; 1 and 2 days after the removal of the inhibitor. Transiently blocking PI3K signaling therefore drives Notch signaling and HES1 protein production throughout organoid nephrons, while control organoid nephrons exhibited low and non-uniform HES1 protein levels (**Figure S3b**). Importantly, inhibition of Ly29 altered the structural dynamics of the organoids, leading to the emergence of JAG1^+^/HNF1B^+^ structures distributed across both the periphery and center of the organoid discs, while control organoids predominantly exhibited tubulogenesis biased towards the periphery (**Figure 2a; Figure S3b**).

**Figure 2.**
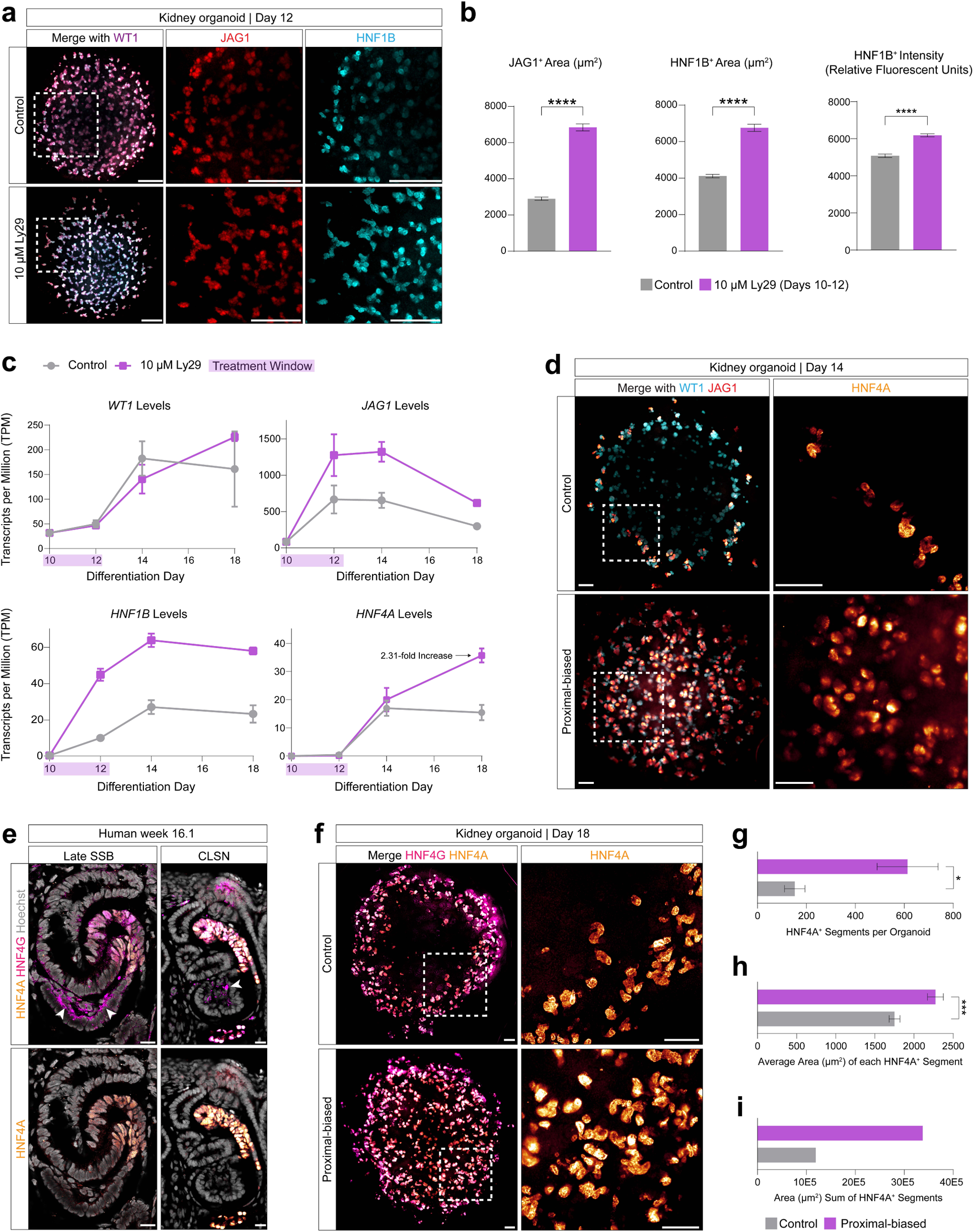
PI3K inhibitor-treated kidney organoid nephrons exhibit a shift toward proximal cell fates. **a**. Whole-mount immunofluorescent stain of day 12 control and LY294002-treated kidney organoids. Insets highlight JAG1 and HNF1B protein expression. Scale bars: 500 microns. **b**. Measurements from (a) for JAG1^+^ and HNF1B^+^ nephron size (µm^2^) and HNF1B^+^ intensity (relative fluorescent units) for *n* = 3 day 12 organoids. SEM error bars are shown. Statistical significance is determined using Student’s t test. **c**. Bulk RNA-sequencing (TPM) of select genes on days 10, 12, 14, and 18. SEM error bars are shown from *n* = 2 whole organoids. **d**. Whole-mount immunofluorescent stain of control and proximal-biased day 14 organoids with insets showing HNF4A expression. Scale bars: 200 microns. **e**. Immunofluorescent antibody stains of week 16 human kidneys during HNF4A^+^, HNF4G^+^ proximal tubule elongation. White arrowheads indicate autofluorescence from endothelial cells. Scale bars: 10 microns. **f**. Whole-mount immunofluorescent stain of control and proximal-biased day 18 organoids with insets showing HNF4A expression. Scale bars: 200 microns. **g**. Quantification of total number of HNF4A^+^ organoid nephron segments from *n* = 4 whole day 18 organoids. SEM error bars are shown. Statistical significance is determined using Student’s t test. **h**. Quantification of the average area (µm^2^) of HNF4A^+^ organoid nephron segments from *n* = 3 whole day 18 organoids. SEM error bars are shown. Statistical significance is determined using Student’s t test. **i**. Area sum (µm^2^) quantification of HNF4A^+^ organoid nephron segments from *n* = 3 whole day 18 organoids.

*In vivo*, the JAG1^+^/HNF1B^+^ state transitions into HNF4A^+^ proximal cells (**Figure 1b**) with HNF4G co-upregulated when the proximal precursor domain elongates (**Figure 2e**). While control organoids show some HNF4A expression by differentiation day 14 (**Figure 1g**), primarily at the periphery of organoids (**Figure 2d**), PI3K inhibitor-treated organoids upregulated HNF4A extensively throughout organoids (**Figure 2d, 2f**). Treated organoids increased the total number of HNF4A^+^ segments per organoid (4.02-fold increase, *p*-value < 0.05), the average size of individual HNF4A^+^ segments (1.30-fold increase, *p*-value < 0.001), and therefore the sum total area of HNF4A^+^ organoid nephron segments (2.83-fold increase for *n* = 3 organoids) (**Figure 2g-i**). As the organoid nephrons matured, they further upregulated *HNF4A*/HNF4A (2.31-fold increase in transcript levels, *p*-value < 0.05) as well as *HNF4G*/HNF4G (1.72-fold increase in transcript levels, *p*-value < 0.05) (**Figure 2c, 2f; Table S5**). We confirmed this phenotype using a second inhibitor of PI3K signaling that differs structurally from Ly29, GDC-0941^36^ (not shown). Notably, the PB organoids had unaltered *WT1* RNA and protein levels, marking differentiating podocytes (**Figure 2c-d**). The spreading of nephron forming events from periphery to center of each organoid upregulated HNF4A centrally (**Figure S3c-d**). Those nephrons positioned at the periphery increased expression of *HNF4A* and transcription factor *POU3F3*, which is only expressed in segment 3 in the proximal tubule, and otherwise distally. The upregulation of *POU3F3* was confirmed by our bulk RNA-sequencing (**Table S5**). These data show organoid nephrons can be biased to upregulate transcription factors driving early proximal tubule development.

### Enriching a functional proximal precursor identity in kidney organoids

To examine the process by which organoids develop PB cell identities in our protocol, we transcriptionally profiled control and Ly29-treated (PB) organoids using single-cell RNA-sequencing, sampling organoids across time (days 10 [untreated], 12, 14, and 18) and conditions. The organoid separated into nephron-like (*PAX2^+^*) and interstitial-like (*PDGFRA*^+^) cells (**Figure S4a-b; Table S6**). Day 10 organoid nephron cells showed *WT1*^+^/*SIX1*^+^/*PAX2*^+^ progenitor-like profiles (**Figure S4c; Table S6; Table S7**). These cells were positive for *CITED1* and robust expression of *PAX8,* indicating a lack of an *in vivo* nephron progenitor cells where *PAX8* is normally detected at low levels in *CITED1*^+^ cells^37^ (**Table S1**). Organoid nephrogenic cells separated by their sampling time, but overlap was observed between each time, thus forming an inferred developmental trajectory from progenitors through to *HNF4A^+^* cells (**Figure 3a-c; Table S7**). The inferred trajectory suggests day 10 cells transition from progenitors and differentiate into nephron lineages by day 18 as evident from gradual changes in gene expression (**Figure S4d-f**). These data validate our bulk RNA-sequencing and whole-mount immunofluorescent antibody staining, confirming increases in early proximal/medial gene expression such as *HNF1B* under PB conditions, and show elevated *JAG1* precedes *HNF1B* and subsequent *HNF4A* expression (**Figure 3c**). Control and PB organoid nephron cells co-clustered, indicating that early and transient Ly29 treatment coaxed cells towards a PB state, increased the number of these cells, but without eliciting other transcriptional changes that drive a transcriptional state sufficiently distinct to separate clusters. 72.7% of all *HNF4A*^+^ cells originated from the PB condition, with similar ratios in *HNF1B* and *JAG1* (**Figure 3c**).

**Figure 3.**
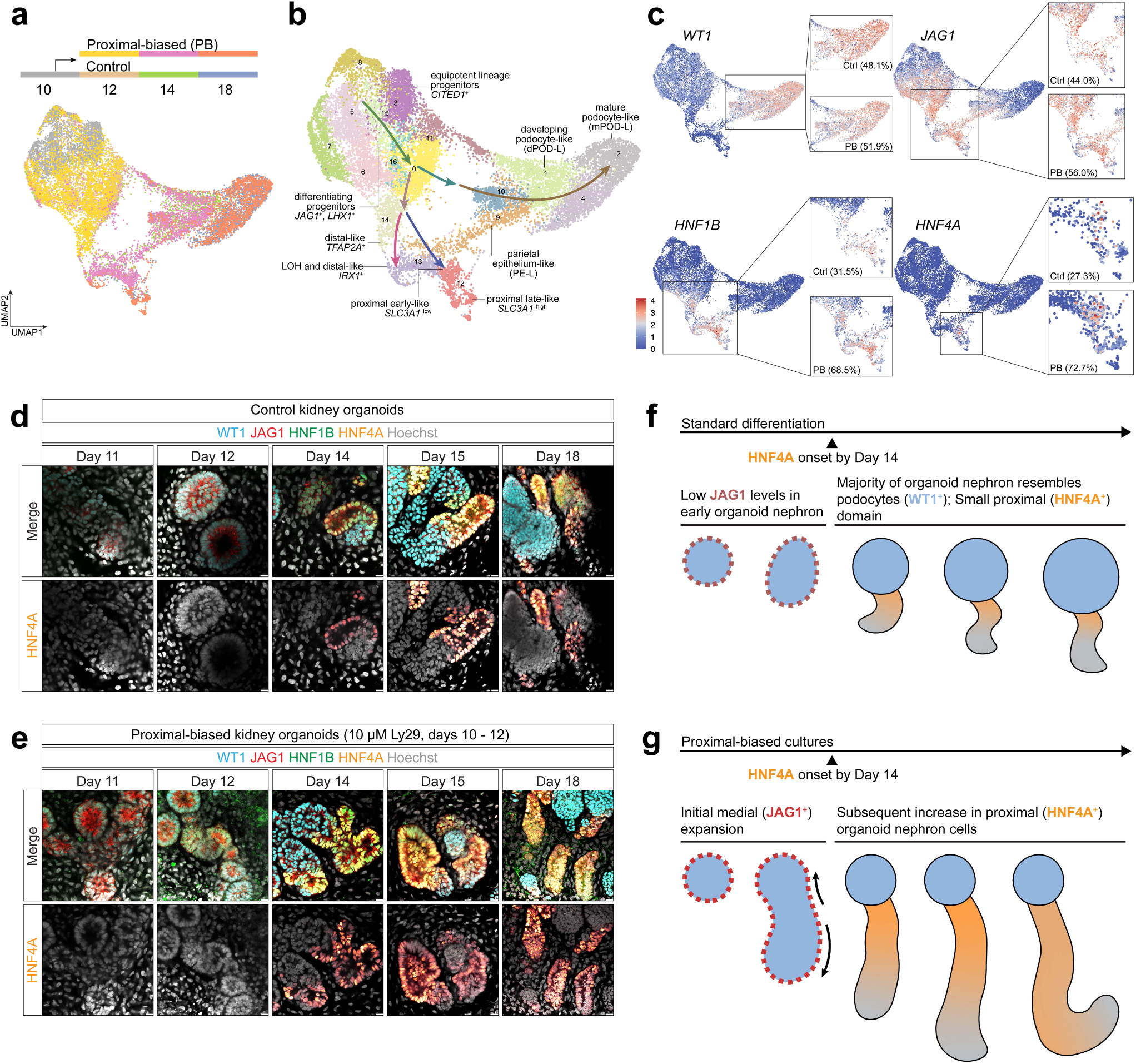
Single-cell transcriptomic landscape and morphological changes in proximal-biased kidney organoids across differentiation stages. **a**. Schematic of single-cell RNA-sequencing timepoints with UMAP reduction of kidney organoid nephrogenic cells, colored by samples. **b**. UMAP reduction of nephrogenic cell clusters in kidney organoids, colored by clusters. **c**. Feature plots for select nephron marker genes with insets separating control from proximal-biased cells, and percent quantification of total cells expressing the gene. **d**. Whole-mount immunofluorescent stains of control kidney organoid nephrons from differentiation days 11, 12, 14, 15, and 18 with the bottom row highlighting HNF4A protein expression. Scale bars: 10 microns. **e**. Whole-mount immunofluorescent stains of proximal-biased kidney organoid nephrons from the same differentiation days as (d), showing HNF4A protein expression in the bottom row. Scale bars: 10 microns. **f**. Differentiation model for control kidney organoids based on (d). **g**. Differentiation model for proximal-biased kidney organoids derived from (e).

To understand the cellular composition of our control and PB organoid nephron cells, we integrated and compared their transcriptional profiles with *in vivo* nephron cells (**Figure S1a; Figure S4g; Table S8**). As expected, organoid nephrogenic cells mostly co-clustered with human nephron cells (**Figure S4h**). Sample contributions for each cluster were quantified (**Figure S4h-i**), revealing parallels and discrepancies between *in vivo* and *in vitro*-derived PB organoid nephron cells. Organoid podocyte (clusters 2 and 3: *MAFB*^+^, *OLFM3*^+^, *NPHS2*^+^), early proximal (cluster 12: *HNF1B*^+^, *JAG1*^high^), proximal (cluster 10: *HNF1B*^+^, *HNF4A*^+^, *SLC3A1*^+^), and loop of Henle precursor (cluster 9: *SLC12A1*^+^, *MAL*^+^) cells co-clustered with *in vivo* cells. In contrast, human *CITED1^+^* nephron progenitors (clusters 0 and 5) did not co-cluster with day 10 organoid cells (cluster 1) as expected since *CITED1*^+^ day 10 organoid cells also express *PAX8* and *LHX1* (**Figure S4g-j; Tables S6, S7, and S8**). Distinct cluster were observed between day 12 control (clusters 7 and 8: *PDGFRB*^+^, PI3K^HIGH^), day 12 PB (cluster 4: *JAG1*^+^, *DLL1*^+^, Notch^HIGH^), and early human nephron (cluster 6: *SNAI2*^+^, *DAPL1*^+^) cells, pointing to transcriptional differences induced by Ly29 treatment (**Figure S4h; Figure S1h-j; Table S8**). Distal human nephron cells (clusters 20, 19, and 16) separated from organoid nephron cells. These data collectively indicate that the PB organoids primarily generate podocyte and proximal cells, and few other cellular identities.

Validating these data in individual organoid nephrons showed consistent views (**Figures 3d-g**). Morphologically, PB organoids diverge from controls, with a shift to HNF4A^+^ cells, leading to an expansion of the HNF1B^+^/HNF4A^+^ identity throughout the tubular portion of the organoid nephron. Initially, PB organoids display a subtle increase in the WT1^+^ domain immediately following PI3K inhibition at day 12 (1.08-fold larger, *p*-value < 0.01) (**Figure S4k-l**). However, this effect is not sustained and compared to control organoids, each control nephron displays large WT1^+^ renal corpuscle-like structures which are larger than those in the PB organoids (1.42-fold larger, *p*-value < 0.001) (**Figure 3d,f; Figure S4k-l**). Instead, the PB organoid nephrons show an elongated HNF4A^+^ proximal domain (**Figure 2f-i**; **Figure 3e,g**) compared to controls (**Figure 3d,f**).

To scrutinize how PB nephrogenic cells compare against other kidney organoid proximal tubule models, we performed a three-way analysis between our PB organoid nephrons (**Figure 3a-c**), cells from a study generating proximal tubule-enhanced kidney organoids^10^ (**Figure S5a-c; Table S9**), and we used the *in vivo* human proximal developmental program as a benchmark (**Figure S1a-c; Table S1**). Because cells from these data were captured using different single-cell technologies, we mapped each organoid dataset to the *in vivo* reference using the Mutual Nearest Neighbor (MNN) method to mitigate batch effects (**Figure S5d**)^38^. In addition, since organoids were sampled at discrete timepoints (**Figure S5e**), while human *in vivo* data represents a continuum of differentiation, we used the previously identified *HNF1B* and *HNF4A*-correlating genes to build a reference for the ‘*early*’ and ‘*late*’ proximal developmental programs (*early*: genes expressed in human clusters 21, 24, and 20; *late*: genes not expressed in human cluster 21, but expressed in clusters 24 and 20) (**Figure S2a**).

Using these reference points, organoid cell profiles were assessed using the distribution of expression of *early* and *late* genes using a Wilcoxon Rank-sum test, incorporating a Bonferroni correction for multiple testing. Early proximal genes in PB organoid nephron cells closely resembled the distribution and expression levels detected in human cells (for instance *ACSM2A*, *SLC39A5*, *UGT3A1*) while proximal tubule-enhanced cells show a distinct grouped expression profile (**Figure S5f**). Among the early proximal genes analyzed, 14/15 displayed an expression distribution statistically closer between the *in vivo* and PB organoid data compared to the proximal tubule-enhanced data (**Table S10**). Similarly, late proximal tubule genes (for instance *OAT3/SLC22A8*, *SLC34A1*, *SERPINA1*) were expressed at lower levels in the *in vivo* and PB organoid cells compared to proximal tubule-enhanced organoid cells (**Figure S5g**; **Table S11**). A likely explanation for these differences is that proximal tubule-enhanced organoid nephron cells reflect a single late timepoint, while PB nephron and *in vivo* cells capture a differentiating proximal program comprising series of proximal precursor states.

### Modeling cisplatin-induced injury and proximal tubule physiologies in kidney organoids

Kidney organoid physiologies are often assessed by cells’ ability to absorb and accumulate compounds that are known to be selectively transported by solute carriers *in vivo*. Uptake of dextran and albumin are associated with the function of HNF4A-target genes *LRP2* and *CUBN*^16,39–42^. PB nephrons display increased expression of *LRP2* and *CUBN* compared to controls (*LRP2*: 1.32-fold higher TPM; *CUBN*: 1.48-fold higher TPM; **Table S5**), raising the possibility that they exhibit proximal tubule functions. To assess this in live cells and to compare to previous kidney organoid models, we used an *HNF4A-YFP* reporter iPSC line to label proximal tubule cells with yellow fluorescent protein (YFP)^43^, and tested whether PB kidney organoids display increased dextran and albumin uptake. Day 18 PB organoids showed strong HNF4A-reporter activity (**Figure S6a**) and increased uptake of fluorescently tagged dextran (**Figure 4a**; 6.87-fold increase, *p*-value < 0.05) and albumin (**Figure S6b**; 4.06-fold increase, *p*-value < 0.05) compared to controls. Albumin and dextran were only detected within HNF4A-YFP^+^ regions of nephrons, and immunostaining post-assay confirmed that uptake was confined to HNF4A^+^, LRP2^+^ nephron segments highlighting proximal precursor-specific roles (**Figure 4b**).

**Figure 4.**
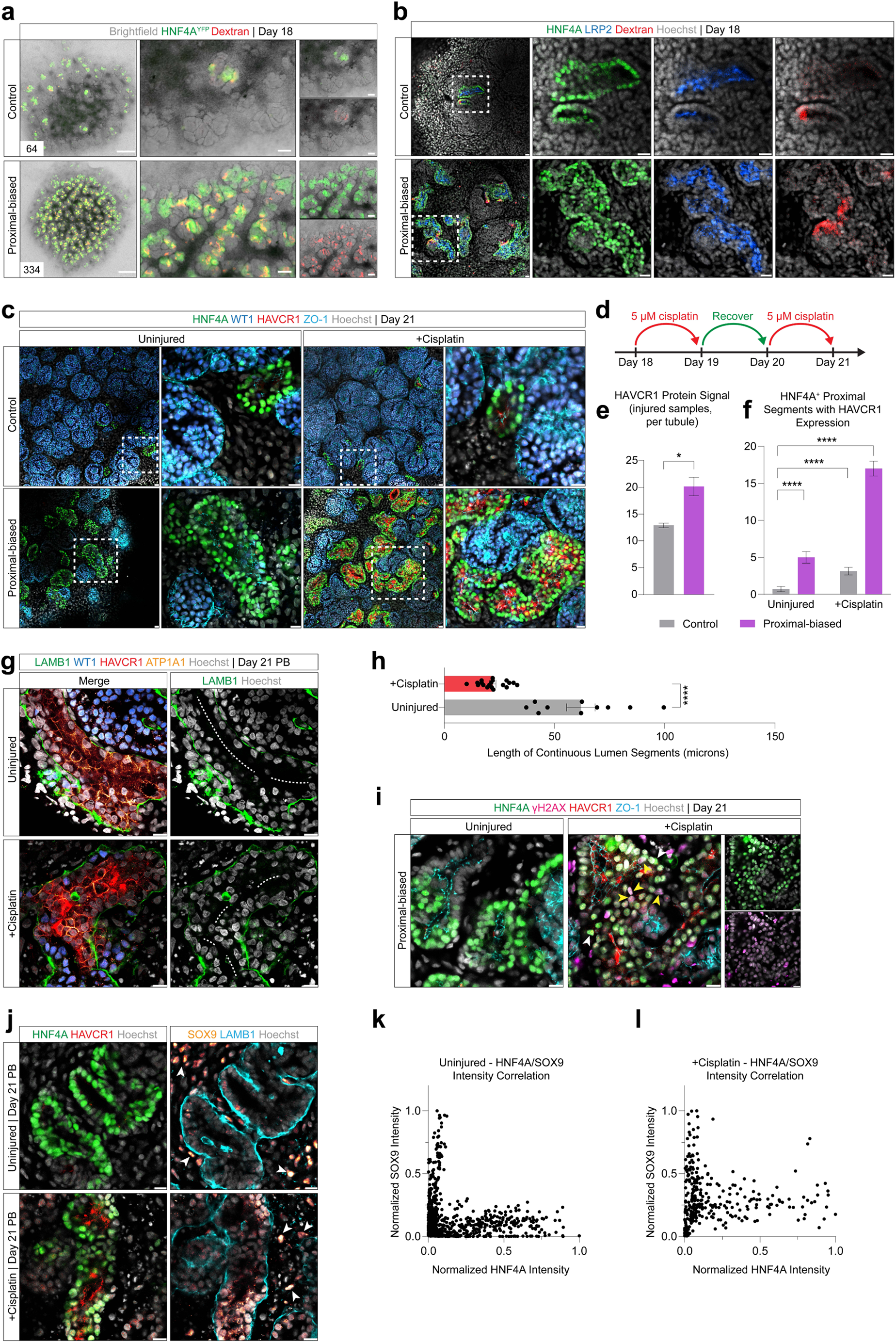
Functional characterization and injury response in proximal-biased kidney organoids. **a**. Live imaging of HNF4A-YFP (Vanslambrouck et al., 2019 *JASN*) kidney organoids on differentiation day 18 with Alexa 647-conjugated dextran uptake. Quantification of individual uptake events displayed in lower left, and insets detail HNF4A-YFP and dextran distinction. Scale bars: whole organoid-500 microns, insets-100 microns. **b**. Whole-mount immunofluorescence of day 18 control and proximal-biased kidney organoids following dextran uptake assay. Boxed region is magnified and split by individual channels. Scale bars: 10 microns. **c**. Whole-mount immunofluorescence of uninjured and 5 µM cisplatin-injured kidney organoids on differentiation day 21, split by control and proximal-biased conditions. Boxed area magnifies HNF4A^+^ proximal organoid nephron regions. Scale bars: 10 microns. **d**. Timeline schematic of kidney organoid injury strategy. **e**. Quantification of relative protein signal from HAVCR1^+^ regions in control and proximal-biased kidney organoid nephrons on day 21 following injury. Data are averaged from *n* = 3 independent and consistent imaging panels. SEM error bars are shown. Statistical significance is determined using Student’s t test. **f**. Quantification of HNF4A^+^ proximal organoid nephron segments with HAVCR1 expression for each of the four conditions from (c). Positive segments are counted from a 20X widefield panel of 4 images per organoid, from *n* = 2 biological replicate organoids for each condition. SEM error bars are shown. Statistical significance is determined using Student’s t test. **g**. Whole-mount immunofluorescence of uninjured and 5 µM cisplatin-injured kidney organoid nephrons on day 21. Dashed white line indicates an uninterrupted segment through the lumen of the organoid tubule. Scale bars: 10 microns. **h**. Quantification of continuous lumen segments from (h). Data are quantified from *n*= 4 organoids per condition, and SEM error bars are shown. Statistical significance is determined using Student’s t test. **i**. Whole-mount immunofluorescence of uninjured and 5 µM cisplatin-injured proximal-biased kidney organoid nephrons on day 21. Yellow arrows denote cells of the HNF4A^+^ organoid nephron tubule with low HNF4A protein expression and positive expression of γH2AX. White arrows indicate cells with high HNF4A, and low γH2AX detection. Split channels separate HNF4A and γH2AX for cisplatin-injured condition. Scale bars: 10 microns. **j**. Whole-mount immunofluorescence of uninjured and 5 µM cisplatin-injured proximal-biased kidney organoid nephrons on day 21. White arrowheads indicate interstitial cells surrounding the tubules that are positive for SOX9. Scale bars: 10 microns. **k-l**. Quantification of HNF4A and SOX9 signal intensities from (j) for uninjured (k) and +cisplatin (l) samples. Intensities were quantified from individual cells of technical and biological duplicates (4 organoids per condition) and normalized using min-max normalization.

Single-cell RNA-sequencing data show PB cells express a broad range of genes associated with proximal tubule physiological functions, such as sodium bicarbonate transport (*SLC4A4*, 71.5% of cells from PB), monocarboxylate and iodide transport (*SLC5A8*, 79.0% of cells from PB), amino acid transport (*SLC3A1*, 71.8% of cells from PB), and copper transport (*SLC31A1*, 71.0% cells from PB) (**Figure S6c**). *SLC31A1* has also been shown to be important for uptake of cisplatin, a chemotherapeutic with severe nephrotoxic side-effects^44–54^. Recent studies show that organoids are sensitive to cisplatin and respond by activating early proximal tubule injury marker *KIM1/HAVCR1* expression and display evidence of DNA damage^12–14^. To determine whether PB cells can model cisplatin-induced injury, we treated PB organoids and controls with two staggered low doses of cisplatin (5 µM) and assayed organoids 3 days after initial injury (**Figure 4d**). Cisplatin treatment resulted in strong detection of KIM1/HAVCR1 in HNF4A^+^ PB nephron cells, while injured control organoids exhibited sparse KIM1/HAVCR1 (**Figure 4c**), with a marked shift of KIM1/HAVCR1 protein abundance in injured PB organoid nephrons (**Figure 4e**, 1.56-fold higher, *p*-value < 0.05). Injured PB organoids contain more KIM1/HAVCR1^+^ and HNF4A^+^ double-positive organoid nephrons compared to injured control organoids, as well as uninjured control and PB organoids (all *p*-values < 0.0001) (**Figure 4f**). Repeat cisplatin-induced injury can drive tubular atrophy in kidney organoids^13^. To determine if just two doses of early cisplatin treatment collapse apical-basal polarities in PB organoids, we analyzed uninjured and cisplatin-treated PB nephron tubules for apical and basal cell surface markers ATP1A1 and LAMB1, respectively (**Figure 4g**). Luminal integrity was lost in injured HAVCR1^+^ organoids, as evidenced by shorter open lumen surfaces, while uninjured controls preserved tubular architecture and maintained continuous open lumens (**Figure 4h**; 2.82-fold longer uninterrupted tubular lumens, *p*-value < 0.0001).

Reports indicate that *Hnf4a/HNF4A* are downregulated in proximal tubules following injury^55–57^ and injury programs are mosaic^58^. To determine whether the varied HNF4A protein levels in injured nephrons suggest different cellular responses, we stained for DNA damage response marker γH2AX^12–14^. Detection of γH2AX coincides with varied HNF4A protein levels, indicating a mosaic response (**Figure 4i**). Of note, interstitial γH2AX was also detected, suggesting broad DNA damage throughout organoids as previously reported^14^. In mice, proximal tubular cells respond to injury by activating *Sox9*^58–61^, and under conditions of ischemia-reperfusion injury and rhabdomyolysis-induced acute kidney injury and disease models, the status of Sox9 ultimately determines proximal tubule regeneration with or without fibrosis^58^. To test if PB kidney organoid nephrons activate SOX9 in response to cisplatin-induced injury, we analyzed uninjured and cisplatin-treated tubules for the coexpression of HNF4A and SOX9. HNF4A and SOX9 exhibited limited coexpression in the uninjured condition, while cisplatin-treated PB organoids activated SOX9 within HNF4A^+^ tubules (**Figure 4j**). Given the mosaic injury response, we quantified SOX9 signal at a single-cell resolution. HNF4A^+^ cells exhibited higher levels of SOX9 signal intensity in the injured condition, indicating activation of SOX9 in organoid proximal cells in response to cisplatin treatment (**Figure 4k-l**). These data show that PB nephron cells provide a sensitive and rapid system to study cisplatin-induced cellular injury responses.

Given the improved capability of PB organoid nephrons to replicate physiological processes and display markers of injury, coupled with the reproducibility of biasing other iPSC lines (**Figure S6d**), this approach emerges as a robust method for generating *in vivo*-mimicking proximal precursor cells within nephrons cultured *in vitro*.

## DISCUSSION

We developed informed approaches to mimic human proximal nephron precursor cells in organoids by delineating early *in vivo* human proximal tubule development, comparing it to existing kidney organoid models, and thereafter driving changes in cell states to bias differentiation outcomes. Our new model generates nephron-like structures in thin self-organizing discs that preserve nephron 3D complexities. We take advantage of the known plasticity in nephron positional identities^25^ and drive organoid nephrons towards a proximal-biased (PB) precursor-like state where organoid cells sequentially activate transcription factors and express *HNF4A* and function-imparting proximal tubule genes. This system offers a reproducible model to study development, origins of congenital disease, kidney injury, and physiology and is compatible with high-resolution microscopy. Our approach is aligned with the broader body of work in kidney organoids, development, and disease modeling, and provides a targeted technique generating proximal tubule cell types with improved fidelity to their *in vivo* counterparts.

### Benchmarking and recapitulating the proximal developmental program

We demonstrate that *in vivo*, human proximal precursors develop through a series of cell-state transitions marked by activation of JAG1, followed by HNF1B, and eventually production of HNF4A in the upper bend of the medial S-shaped body nephron. This is consistent with our previous studies showing human nephrons activate HNF4A in the medial nephron segment^18^, and work by others in the mouse model^23^. Genetic evidence from mice shows that Notch signaling is required for proximal nephron differentiation, nephron segmentation, and nephron number^22,32,33,62–64^. The emerging view is that the requirement for Notch signaling in proximal tubule development is deeply conserved between human and mice.

We further show that inhibition of PI3K results in a transient upregulation of Notch target genes, and upregulation of HNF1B. Inhibiting PI3K and simultaneously blocking Notch signaling in PB organoids prevents HNF1B expression, which is consistent with *Notch1*; *Notch2* double knockout mice failing to upregulate *Hnf1b* in nephrons^22^. *In vivo*, Hnf1b binds to and is required for expression of *Hnf4a*^23^. A conserved role for HNF1B is indicated by mutations in *Hnf1b/HNF1B* being linked to congenital kidney anomalies affecting tubulogenesis, as well as studies in organoids^24,65–69^. In our model, increased HNF1B expression and protein levels precede detection of *HNF4A,* concurring with *in vivo* and *in vitro* studies showing HNF4A is essential for proximal nephron cell-fate development^15–17^.

While *Hnf4a*/*HNF4A* and *Hnf4g*/*HNF4G* are co-expressed in proximal tubules *in vivo*^8,18,27^, evidence suggests a non-redundant relationship. *Hnf4a* knockout mice and *HNF4A* knockout kidney organoids have prominent phenotypes^15–17^. Conversely, loss of *HNF4G* moderately alters kidney organoids^15^, and *Hnf4g* knockout mice display a phenotype in the intestine only when removed alongside *Hnf4a*, due to the redundant roles of Hnf4 factors in enterocyte maturation^70,71^. As part of our comparative analyzes between *in vivo* and *in vitro,* we sought to understand how Hnf4a-dependent genes are expressed during development and in adult nephrons. It is interesting to note that a group of Hnf4a-dependent genes are not expressed in the adult, indicating temporally dynamic roles for Hnf4a, or that gene expression is mediated by indirect mechanisms requiring for example co-factors. Given the clear role of Hnf4a in injury and repair mechanisms, this is an area needing further scrutiny. Overall, our analyses point to that human orthologs of Hnf4a-dependent genes are expressed at much lower frequencies in organoids, which in turn suggests that organoids at present only partially mirror the *in vivo* proximal program. However, the onset of HNF4A and HNF4G expression occurs within a short timeframe in the S-shaped body nephron and PB organoids do express both genes as the nephron tubules elongate, which mirrors the temporal *in vivo* sequence. Other kidney models generating HNF4A^+^ proximal cell-states have previously enriched kidney organoid cultures for the nephron progenitor state and subsequently optimized culture conditions to permit proximal-like cell development^10^. In contrast, the PB system imposes a specific signaling event switching cells to a proximal developmental program. For improving reproducibility, synchronicity, and application across cell-lines, increasing control over differentiation outcomes is an efficient strategy.

In all kidney organoid systems, it is unclear how JAG1 is initially activated in the forming nephron. *In vivo,* it is thought that *Jag1* is bound and sensitive to β-catenin mediated transcription^72,73^ and Jag1/JAG1 expression initiates in cells adjacent to the WNT9B secreting collecting duct^21,62,74^. It is unclear how the brief dosing with CHIR at day 7 (a GSK3-β antagonist and WNT/β-catenin agonist) can control the emergence of JAG1^+^ nephron aggregates at day 10. Moreover, *in vivo* JAG1 expression follows an intriguing wave-like pattern where it moves proximally through the nephron and is downregulated distally, suggesting multiple layers of control over this Notch ligand^21^. Identifying the mechanisms that regulate *JAG1* expression and activities will be critical for understanding nephron proximal-distal patterning.

### Modeling physiologies and injury in proximal-biased nephrons

*In vivo*, proximal tubule cells exhibit a well-defined apical-basal polarity, appropriately positioned solute carriers and transporters, vascularization, susceptibility to intraluminal solute flow, and sensitivity to nephrotoxic compounds^1,75–77^. Modeling nephrotoxicity in PB organoids therefore lacks flow and vascularization, but they develop an apical-basal polarity and accumulate dextran and albumin in HNF4A^+^, LRP2^+^ cells. We show that PB nephrons are sensitive to Cisplatin early in the differentiation protocol (day 18), at a low concentration (5µM), and within a quick timeframe (72 hrs), making the model suitable for screening experiments requiring short dosing plans and quick endpoints. Cisplatin is transported by a range of solute carriers^44,78,79^, and bulk RNA-sequencing and single-cell RNA-sequencing confirm expression of several (maximum TPM values *CTR1/SLC31A1*: 120.92; *OAT1/SLC22A6*: 2.89; *OAT3/SLC22A8*: 3.83; *OCT2/SLC22A2*: 3.89) (**Table S5**). However, while detected, it remains to be determined whether cisplatin-induced damage requires active transport. Expression of *HAVCR1/KIM1* was specific to HNF4A^+^ cells, as would be expected but DNA-damage marker γH2AX was detected throughout organoids, corroborating recent findings^14^. Dissecting the mechanism by which tubular HNF4A^+^ cells are injured by cisplatin will further validate its use as a proximal tubule nephrotoxicity model. We show that HNF4A^+^ organoid nephrons upregulate SOX9 in response to cisplatin. Recent studies have demonstrated organoids possess intrinsic repair capabilities but this repair capacity is diminished following repeated cisplatin exposure, partly due to tubular atrophy^13^. Our results indicated that proximal-biased kidney organoid nephrons undergo tubular collapse after just two low doses of cisplatin. Future experiments are required to assess the regenerative potential of these organoids under prolonged culture conditions and to determine the timeline and extent of possible recovery following repeated cisplatin treatment. Developing a tightly controlled synchronized injury model suitable for posing precise questions on the impact of compounds – such as chemotherapeutics, antibiotics, or other agents – is of high clinical value and would allow scrutiny of transition states that proximal nephron cells undergo post-injury^80^.

### Model limitations and integration with other systems

The current PB model represents a cell-state that we best estimate to resemble a capillary loop stage based on the expression of some solute carriers but lacking a full complement of mature nephron markers. Generating a mature proximal tubule will require further modifications including tuning of culture conditions to drive maturation, elongation, and separation of proximally distinct segments. Beyond manipulation of pathways, the PB model lacks physiological properties such as flow, vascularization, and appropriate culture conditions including supportive interstitial cell-types. Evidence shows that sorted proximal tubule-like cells from hiPSC-derived kidney organoids grown with or without immortalized cells can recapitulate proximal tubule physiologies^81,82^ and isolated proximal nephron tubules can be cultured for months, which introduces a direct possibility to adopt our protocol and generate PB organoids that are used as a cell source for long-term tubule culture systems. Organoid production can also be scaled to be suitable for large-scale screens, as shown in experiments using a bioreactor to produce kidney organoids^69^. On their own, these organoids did not exhibit extensive proximal tubule development but do generate some LRP2^+^/Megalin^+^ structures, again raising the possibility of applying our approach to bias them to a proximal lineage. Beyond the manipulation of culture methods in static or dynamic systems, proximal nephron cells isolated from kidney organoids have been used in bioengineered chips to study pharmacological drug uptake^83^. Chip technologies have further allowed for the introduction of nephrons with vasculature to model renal physiologies^84^. These methods and the rapidly developing kidney organoid field contribute to the expanding toolkit available for kidney research and complement our developmental-mimicry approach.

## METHODS

### Human kidney samples

Human fetal samples were collected under Institutional Review Board approved protocols (USC-HS-13-0399 and CHLA-14-2211). Following the patient decision for pregnancy termination, informed consent for donation of the products of conception for research purposes was obtained and samples collected without patient identifiers. The person obtaining informed consent was different than the physician performing the pregnancy termination procedure, and the decision to donate tissue did not impact the method of pregnancy termination. Fetal age was determined according to the American College of Obstetrics and Gynecology guidelines. The kidney samples ranged from 14 to 17 weeks of gestation with no sex reported. Intact samples within the kidney capsule were analyzed. Samples were transported on ice at 4°C in high glucose DMEM (Gibco, 11965-118) supplemented with 10% fetal bovine serum (Genesee Scientific, 25-550) and 25mM HEPES (Gibco, 15630080).

### Human induced pluripotent stem cell lines

An iPSC line (male) was provided by Dr. Amy Firth (USC). An HNF4A-YFP reporter iPSC line (male) was obtained through the Washington University Kidney Translational Research Center (KTRC) and the ReBuilding a Kidney (RBK) Consortium. Details on the *HNF4A-YFP* iPSC line can be found elsewhere^43,85^.

### Mouse kidney explants and culture

Pregnant female Swiss Webster mice were euthanized on day E12.5 of pregnancy via CO_2_ following appropriate University of Southern California IACUC approved protocol. Embryos were removed and placed into PBS before dissecting out kidneys for explant culture as described below.

### iPSC maintenance

*Matrigel-coated plate preparation*. DMEM (Corning, 10-017-CV) was aliquoted into a 15 mL conical vial, and 120 µL of Matrigel (Corning, 354277) was added to make a 1% Matrigel mix. After thorough mixing, 2 mL of 1% Matrigel was pipetted into each well of a 6-well plate (or 1 mL/well for a 12-well plate). Matrigel plates were then incubated at 37°C/5% CO_2_ overnight before use.

*Biolaminin 521 LN-coated plate preparation.* DMEM was aliquoted into a 15 mL conical vial and 300 µL of biolaminin 521 LN (from Biolamina, LN521) was added to make a 5% biolaminin 521 LN mix. After thorough mixing, 0.5 mL of 5% biolaminin 521 LN was pipetted into each well of a 12-well plate. Biolaminin 521 LN plates were then incubated at 4°C overnight before use.

*iPSC expansion and maintenance*. iPSCs were thawed in Essential 8 media (Thermo Fisher Scientific, A1517001) on 1% Matrigel-coated plates. Media was initially supplemented with 10 µM Y-27632 (Rho kinase inhibitor from Tocris, 1254) for 24 hours after thawing iPSCs. Media was changed every 24 hours until cells reached 70-80% confluency (3 days). For freezing, when iPSCs reached 70-80% confluency, cells were collected for passage as described below and resuspended into a mix of 50% Essential 8, 40% KnockOut Serum Replacement (Thermo Fisher Scientific, 10828010), and 10% DMSO (Millipore Sigma, D5879). Cells were then aliquoted into 2 mL cryovials at 500,000 cells/vial, gradually frozen at-80°C overnight, and then transferred to liquid nitrogen storage.

### Directed differentiation to generate kidney organoids

Differentiation protocols were developed based on published protocols^86,87^ and adapted in our laboratory. Each biological replicate was generated from a distinct frozen vial of iPSCs. On reaching 70-80% confluency (3 days), iPSCs were rinsed with 1xPBS (Thermo Fisher Scientific, 10010049) and then incubated in TrypLE Select Enzyme (Thermo Fisher Scientific, 12563011) for 6 minutes at 37°C/5% CO2. The enzymatic reaction was neutralized using a volume of Essential 8 media 2X that of TrypLE. Cells were collected and resuspended in Essential 8 media supplemented with 10 µM Y-27632, with 10,000 cells/well plated onto a 5% biolaminin 521 LN-coated 12-well plate. 6 hours after plating, differentiation was initiated by changing culture media to TeSR-E6 (Stem Cell Technologies, 05946) supplemented with CHIR99021 (Tocris, 4423). Briefly, culture medium was supplemented with CHIR99021 for 5 days (exact concentration and duration are dependent on cell line being used), followed by 2 days with 200 ng/mL FGF-9 (R&D Systems, 273-F9) and 1 µg/mL Heparin (Millipore Sigma, H4784). At day 7, the cells were detached using TrypLE as described above and resuspended in TeSR-E6 supplemented with 10 µM Y-27632. 200,000 cells were seeded into each well of a round-bottom non-adhesive 96-well plate. Organoids were manually transferred to a 0.4 µm pore culture plate (Corning, 3450; Stem Cell Technologies, 100-1026). Organoids received a pulse of TeSR-E6 supplemented with CHIR99021 for 1 hour; media was then switched back to TeSR-E6 supplemented with 200 ng/mL FGF-9 and 1 µg/mL Heparin until day 12. Proximal-biased kidney organoids were cultured in TeSR-E6 supplemented with 200 ng/mL FGF-9, 1 µg/mL Heparin, and 10 µM LY294002 (Tocris, 1130) for 48 hours between differentiation days 10 and 12; untreated control organoids were cultured with 200 ng/mL FGF-9, 1 µg/mL Heparin, and DMSO vehicle. From day 12 onward, organoids were cultured in TeSR-E6 alone.

### Small-molecule inhibitor screen and culture optimization

For all small molecule inhibitors used in this study, initial screens were performed on kidney organoids to determine the optimal dosage and duration of culture. To determine optimal dosage of a molecule, kidney organoids were cultured for 48 hours between differentiation days 10 and 12 in TeSR-E6 supplemented with the molecule at varying concentrations. Concentrations tested used insights from prior studies^25,29^. After an ideal concentration was established, to determine optimal duration of treatment with the small molecule, separate differentiation experiments were set up with the small molecule from differentiation days 10 to 18, as well as differentiation days 12 to 18. For all optimization screens, time-course bright-field imaging was performed, and organoids were collected for whole-mount immunofluorescent analyses on differentiation day 18.

### Mouse kidney organ culture

Kidneys were dissected from E12.5 embryos and cultured for 48 hours at 37°C/5% CO2 in DMEM. Treatment condition culture media was initially supplemented with 20 µM LY294002, 2 µM DAPT (Tocris 2634), or both, before 48 hours of culture, following prior studies^25,29^. After culture, mouse kidney explants were prepared for immunofluorescent analyses as described below for organoids.

### Kidney organoid cell dissociation

Approximately 6 organoids from multiple independent wells of both untreated and treated (48 hour culture with 10 µM LY294002 between days 10 and 12) conditions were collected at differentiation days 10, 12, 14, and 18 for single-cell RNA-sequencing, from separate differentiation batches. The organoids were dissociated using Accumax (Stem Cell Technologies, 07921) at 37°C/5% CO2 for 35 minutes and pipetted twenty times with a P-1000 wide-bore pipette tip every 5 minutes. The dissociation was neutralized by using a 2X volume of AutoMACS Running Buffer (Miltenyi Biotec, 130-091-221). Cells were pelleted by centrifugation at 4°C for 3 minutes at 300 x g. After resuspending the pellet in 2 mL of AutoMACS Running Buffer, cell suspensions were strained through 40 µm strainer (VWR, 21008-949), washing the filter with an additional 1 mL AutoMACS. The flow-through was pelleted, again at 300 x g in a 4°C for 3 minutes. Cells were gently resuspended in AutoMACS Running Buffer supplemented with 14 µM DAPI (Thermo Fisher Scientific, D1306) and 5 µM DR (Thermo Fisher Scientific, 62-254), and sorted on an ARIA II FACS at a low flow rate. Viable cells (DR-positive, DAPI-negative) were collected into a 1.7 mL low binding tube (Corning, 3207).

### SPLiT-seq single-cell RNA-sequencing data collection and analyses

*SPLiT-seq cell processing.* Sorted, single, live cells were immediately processed and fixed using Evercode Cell Fixation kit (Parse Biosciences, ECF2001). Fixed cells were then stored at-80°C until all samples were ready to be processed with the Evercode WT v2 single-cell RNA-sequencing platform (Parse Biosciences, ECW02030). After recovery of barcoded cells, cDNA was cleaned up and amplified by PCR, as per Evercode WT v2 platform protocols. cDNA quality was examined at multiple points on a 4200 TapeStation (Agilent) for yield and quality assessment. Paired-end sequencing was performed on the NovaSeq X Plus PE150 (Novogene).

*scRNA-seq analyses.* From fastq files, demultiplexing, quality control, alignment to reference genome (hg38 Ensembl 105 annotation), and generation of count tables were done using *splitpipe* (Parse Biosciences). The Seurat 4.0 package was used for single-cell RNA-sequencing (scRNA-seq) analyses^88^. To filter out low-quality cells, we kept cells with between 500 and 13000 features, between 100 and 200000 RNA counts, and less than 35% mitochondrial gene content. Sample datasets were integrated using Seurat integration functions across organoid time-points and experimental variables^89^. 50 PCs were used to calculate cell relationships, cluster assignment, and UMAPs. After quality control and sample integration, 56,320 cells remained. Differentially expressed genes of each cluster were identified using the *FindAllMarkers* function, selectively returning genes with expression in at least 25% of cells within the cluster (*min.pct* = 0.25) and with a minimum fold change of 0.25 (*logfc.threshold* = 0.25), while restricting the output to only positively expressed markers (*only.pos* = TRUE). Nephrogenic cell-clusters were identified based on expression of nephron lineage and nephron cell-fate markers and interstitial lineages similarly separated by their gene enrichment. Nephron-forming cells were selected by serial clustering and subsetting while monitoring cluster-based quality metrics. Clusters 3, 6, 8, 9, 10, 13, 14, 15, 16, 17, 20, 24, 26, and 27 were subset for nephrogenic lineage examination.

In vivo *kidney datasets. In vivo* datasets of human week 14^18^ (GSE139280) and human week 17^8^ (GSE124472) were analyzed independently (Figure S1) and were used for comparison to scRNA-seq profiles of the nephrogenic subset of differentiating organoids. To identify genes that are co-expressed with *HNF1B* and *HNF4A*, a Pearson correlation test was performed for each factor, and visualized against clusters 9 (*JAG1*^+^, *HNF1B*^LOW^, *HNF4A*^-^), 21 (*JAG1*^+^, *HNF1B*^+^, *HNF4A*^LOW^), 24, and 20 (*JAG1*^LOW^, *HNF1B*^+^, *HNF4A*^+^), which served as the early medial to proximalizing lineage (Figures S1a and S2a). The expression of these genes *in vivo* and in corresponding clusters of reanalyzed kidney organoid nephrons (Figure S1d; assembled as described below) was analyzed in Figure S2a. Lastly, this human scRNA-seq dataset was integrated with the proximal-biased organoid dataset (Figure S4g-h) using *RunCCA*^88^.

In vitro *organoid datasets.* To unbiasedly assess transcriptional profiles of proximalizing nephron cells, we assembled and reanalyzed differentiating kidney organoids generated from multiple sources. Data from day 25 (GSE102596)^6^; days 7, 15, and 29 (GSE136314)^7^; days 16 and 28 (GSE124472)^8^; and day 26 (GSE118184)^9^ organoids were reanalyzed. Raw data from each dataset were acquired and analyzed individually as above before being integrated. This integrated dataset was subset into nephrogenic cell types for direct comparison to *in vivo* nephrogenic cells (Figure S1). Here, clusters 7, 6, 14, 12, 8, 18, and 3 served as the proximalizing lineage (Figure S2). Separately, we re-analyzed published scRNA-seq data from organoid nephrons from a protocol generating proximal tubule-enhanced organoids (GSE184928)^10^. These data were acquired and re-analyzed independently. While proximal-biased organoid nephron cells from this study were sampled for control and PB conditions at day 10 (untreated), 12, 14, and 18, representing 3, 5, 7, and 11 days after initiation of 3D organoid culture, single cells from proximal tubule-enhanced organoids were sampled at one timepoint, day 27, representing 13 days of extended monolayer culture followed by 14 days of 3D culture^10^. The *in vivo* cells for this comparison were sampled from a continuum of differentiation (Figure S4e).

*Integration and batch effect correction with MNN*. To analyze the expression patterns of genes associated with proximal tubule program in two organoid datasets where cells were captured using different technologies (Schnell et al., Parse Biosciences SPLiT-seq Evercode WT v2; Vanslambrouck et al., 2022^10^, 10X Genomics Chromium Next GEM Single Cell 3’ Reagent Kits v3.1), we employed a developing human nephron dataset (Figure S1a-c) (Lindström et al., 2021; Tran et al., 2019)^8,18^ as a reference and mapped each organoid dataset to the human reference. To mitigate batch effects, the Mutual Nearest Neighbor (MNN) method was used^38^, selected for its parsimonious correction capability and its proven efficacy in a recent benchmarking study^90^. We retained the shared cell types among the three datasets, specifically early tubules, podocytes, proximal tubules, and distal tubules. These shared cell types facilitated the individual alignment of each organoid dataset with the human developing kidney reference. For the alignment, we focused on the top 3000 highly variable genes with default parameters. To statistically compare the organoid datasets with the human reference data, we conducted a Wilcoxon Rank-sum test, incorporating a Bonferroni correction for multiple testing. The criteria for identifying significantly differentially expressed genes were set at an adjusted *p*-value of less than 0.05, coupled with a log fold change greater than 0.5 or less than-0.5.

*Pseudotime reconstruction of lineages*. Podocytes were subset from the organoid nephron single-cell RNA-sequencing dataset for a more streamlined developmental trajectory analysis from a day 10 equipotent lineage progenitor cell to a day 18 proximal-biased nephron cell. Monocle 3 was used to reconstruct the differentiation trajectory^91–94^.

*Other datasets referenced*. To analyze Hnf4a-dependent gene expression in kidney organoids, we acquired bulk RNA-sequencing data from control and *Hnf4a*-mutant P0 mouse kidneys (GSE144772)^16^. Bulk RNA-sequencing fastq files from two control and two mutant (*Osr2Cre*-driven deletion of *Hnf4a*) P0 kidneys were acquired. Sample quality was assessed using fastqc. Using STAR version 2.7.10a, reads were aligned to the mouse mm10 genome, Ensembl release 102. Mapping quality was appraised using qualimap. Resulting.bam files were then used to generate a feature counts table using *featurecounts*. A list of 257 genes that are significantly downregulated in the *Hnf4a*-mutant kidney (DESeq2 comparing control vs. mutant kidneys, *p*_adj_ < 0.01, log_2_FC ≥ 1.5) were used for analysis^28^ (Figure S2). Separately, a list of 415 genes that are highly expressed and enriched in the adult mouse proximal tubule^27^ (GSE129798; Kidney Cell Explorer; https://cello.shinyapps.io/kidneycellexplorer/; https://github.com/kimpenn/kidneycellexplorer) were used for analyses in Figure S2. To generate the heatmaps presented in Figure 2b-c, three proximalizing human clusters (Figure S1a, clusters 20, 21, and 24) were analyzed, alongside the organoid data, which exclusively focused on cluster 3. These clusters were selected based on their relatively elevated expression levels of *HNF4A*, each surpassing 15% of all cells in the cluster. Human orthologs of mouse genes were retrieved using Ensembl BioMart, and genes expressed in less than three cells of the human dataset were removed.

### Immunofluorescent imaging and analyses

*Kidney organoids and explants.* Whole kidney organoids and mouse kidney explants were fixed for 20 min on ice in 4% PFA (Electron Microscopy Sciences, 15710) in 1xPBS. In 1xPBS, organoids and explants were then carefully cut out of the permeable membrane well and placed into 1xPBS with 1.5% SEA Block (Thermo Fisher Scientific, 37527X3) and 0.1% TritonX100 (EMD Millipore, 1.08643) (blocking solution) at 4°C with gentle movement for one hour. Primary antibodies were resuspended in blocking solution, and organoids were incubated in primary antibody and blocking solution overnight. Samples were rinsed and then washed for at least 3 hours through several rounds of 1xPBS with 0.1% TritonX100. Secondary antibodies were resuspended in blocking solution, and organoids were incubated in secondary antibody and blocking solution overnight. Rinsing and washing steps were repeated the next day before a 25 minute counterstain incubation in 1xPBS supplemented with 0.1% TritonX100 and 1 µg/mL Hoechst 33342 (Thermo Fisher Scientific, H3570) to stain nuclei. Organoids were then washed for 1 hour in 1xPBS alone. Individual organoids were mounted for imaging on slides with a glass coverslip in Immu-Mount (Thermo Fisher Scientific, 9990402). Slides were stored at 4°C in the dark. Primary antibodies used in this study were: WT1 (abcam, ab89901, 1:1000), JAG1 (R&D Systems, AF599, 1:300), HNF1B (Thermo Fisher Scientific, MA5-24605, 1:500), HNF4A (R&D Systems, MAB4605, 1:200), CDH1 (BD Biosciences, 610181, 1:300), ZO-1 (Thermo Fisher Scientific, 33-9100, 1:200), PAX2 (R&D Systems, AF3364, 1:50), SIX1 (Cell Signaling Technology, 12891S, 1:300), HES1 (Cell Signaling Technology, 11988, 1:300), POU3F3 (Novus Biologicals, NBP1-49872, 1:500), HNF4G (Thermo Fisher Scientific, PA5-82189, 1:200), LRP2 (My Bio Source, MBS690201, 1:500), HAVCR1 (R&D Systems, AF1750, 1:200), γH2AX (Cell Signaling Technology, 2577), LAMB1 (Santa Cruz Biotechnology, sc-33709, 1:250), ATP1A1 (Abcam, ab7671, 1:200), and SOX9 (Abcam, ab185230, 1:300). Secondary antibodies conjugated with AlexaFluor 488, 555, 594, and 647 were all diluted to 1:500 in blocking solution.

*Human kidney sections*. Human kidneys were carefully dissected out from donated tissues and placed in 1xPBS. Kidneys were then placed into 4% PFA to fix overnight (18 hrs). Kidneys were washed twice in 1xPBS (1 hour per wash), and samples were placed in 30% sucrose in 1xPBS rocking at 4°C overnight (18 hrs). The next day, kidneys were swirled in Optimal Cutting Temperature (OCT from Sakura Finetech USA Inc, 4583) compound 3 times before embedding in an OCT block. The OCT blocks with tissues were frozen on a dry ice/ethanol slurry. Blocks were stored at-80°C, and ∼12 µm sections were obtained using a cryostat. Slides were stored at-80°C before being processed for immunofluorescent imaging.

### mRNA-sequencing and data analyses

Whole organoid samples (3 independent biological replicates per timepoint, per condition for differentiation days 10 and 12, and 2 replicates for differentiation days 14 and 18) were prepared and purified according to the RNeasy Mini Kit (Qiagen, 74104). Purified RNA was sent to Novogene for sequencing. Reads were aligned to the hg38 Ensembl 105 annotation using STAR version 2.7.10a^95^. A noise-reduction filter was applied to keep genes in which the maximum count in at least one sample was greater than or equal to 10. Partek Flow version 10.0 was then used to normalize these raw feature counts following the transcripts per million (TPM) normalization method. After filtering genes with TPM values greater than or equal to 25 in at least one sample, Z-score averages were calculated and heatmaps were clustered using *pheatmap*. Separately, gene set enrichment analyses were performed specifically on Day 12 replicate samples by first normalizing noise-reduced feature counts using the DESeq2 median ratio method^28^, as recommended for gene set enrichment analyses. Partek Flow version 10.0 was then subsequently used for gene set enrichment analyses.

### *In vitro* TRITC-albumin and dextran uptake assays

For the albumin uptake assay, differentiation day 18 control and proximal-biased organoids were incubated at 37°C/5% CO_2_ for 4 hours in TeSR-E6 supplemented with 10 µg/mL TRITC-albumin (Sigma Aldrich, A2289). Control organoids were incubated with TeSR-E6 alone. After the 4 hour incubation, organoids were washed at least 3 times with TeSR-E6 media and live-imaged immediately after. Organoids were left in culture with only TeSR-E6 media overnight, and then live-imaged again the next day before being fixed and processed for immunofluorescent imaging and analyses. For the dextran uptake assay, organoids were treated and processed in the same way, except that TeSR-E6 was supplemented with Alexa-647-conjugated Dextran (Thermo Fisher Scientific, D22914).

### *In vitro* cisplatin injury assay

For the kidney organoid cisplatin injury assay, we adapted an approach undertaken by two recent studies^13,14^. Differentiation day 18 control and proximal-biased organoids were incubated overnight (18 hrs) at 37°C/5% CO_2_ in TeSR-E6 supplemented with 5 µM cisplatin (Millipore Sigma, 232120). Organoids recovered between days 19 and 20, before another overnight culture (18 hrs) between days 20 and 21 with 5 µM cisplatin. Uninjured organoids were cultured with DMSO vehicle. Organoids were then processed for immunofluorescent imaging and analyses.

### Image acquisition and analysis

Image acquisition of whole-mounted organoids, cryo-sectioned human kidneys, and whole-mounted mouse kidney explants were performed using (1) Leica SP8-X confocal fluorescence imaging system (Leica Microsystems, Germany), in 1024 x 1024 pixels using 25X water, 40X oil, and 63X oil objectives, and (2) Leica Stellaris confocal microscope at 20X oil and 93X glycerol objectives. A Leica Thunder microscope was additionally used at 10X dry objective to image whole-mounted kidney organoids. Live organoids were captured using Zeiss Axio Zoom at 16X and 32X objectives. Image masking was performed and quantified with Imaris 9.7 software (Oxford Instruments), and with Fiji^96^. Fiji was used for individual cell and nephron segment counting, and for quantifying relative protein signal and organoid nephron size.

## ACKNOWLEDGEMENTS

We thank all past and present members of the Lindström Laboratory and extend our gratitude to Dr. Andrew McMahon and Dr. Zhongwei Li for their input as well as several members in their groups. We further thank Dr. Alicia McDonough for insights to human physiology, Dr. Seth Ruffins and the Optical Imaging Facility for assistance with microscopy, and Dr. Bernadette Masinsin and the Flow Cytometry Facility for help with cell isolation. Lastly, we thank Dr. Amy Firth (USC) for providing iPSCs used in this study. J.S. was supported by the T32 Training Grant in Development, Stem Cells, and Regenerative Medicine from NICHD (T32HD060549).

## AUTHOR CONTRIBUTIONS

J.S., and N.O.L. planned experiments and analyzed data. J.S. assembled the figures. J.S., Z.M., M.A., C.C.F., V.W., and F.D.K. performed the experiments, data collection, and/or data analysis. Z.M., J.K., planned and conducted data analyses. M.T. and B.G. provided human tissue. J.S. and N.O.L. wrote the manuscript and incorporated input from all authors.

## CORRESPONDENCE

Further information and requests for resources and reagents should be directed to and will be fulfilled by the lead contact, Nils O. Lindström.

---

Figure S1 Comparative transcriptomics and analyses of in vivo and early in vitro-forming nephrons.

**a**. Single-cell RNA-sequencing of nephrogenic and ureteric lineages from week 14 and 17 developing human kidneys compiled from Lindström et al., 2021 *Dev Cell* and Tran et al., 2019 *Dev Cell*. **b**. Dot plot with identifying genes for each cluster. Arrows, line colors, and cluster numbers match to (a) and (b). **c**. Feature plots for select genes. Insets highlight transition from pretubular aggregate to later nephron stages and specified gene dynamics. **d**. Single-cell RNA-sequencing from four kidney organoid datasets (Combes et al., 2019 *Genome Medicine*; Subramanian et al., 2019 *Nature Communications*; Tran et al., 2019 *Developmental Cell*; Wu et al., 2018 *Cell Stem Cell*). Displayed are only nephrogenic cells ranging between day 7 to day 29 differentiation timepoints. **e**. Dot plot with identifying genes per cluster. **f**. Feature plots for select genes as shown for *in vivo* in (c). **g**. Hierarchical clustering from bulk RNA-sequencing of kidney organoids at differentiation days 10, 12 (control) and 12 (treated for 48 hours with 10 µM LY294002). Samples were averaged from *n* = 3 biological replicates for each timepoint. Clustered genes represent those with a transcripts per million (TPM) value of greater than or equal to 25 in at least one of the samples, (6703 genes). Representative genes for each cluster are shown on the side, along with a Z-score legend. **h-j**. Gene set enrichment analysis plots from Day 12 LY294002-treated / Day 12 control kidney organoids. Plots depict pathway effects from treatment with the PI3K inhibitor LY294002: downregulated (PI3K-Akt signaling pathway, j), upregulated (Notch signaling pathway, k), and unaffected (Hedgehog signaling pathway, l). Normalized enrichment scores (NES), false discovery rates (FDR), and *p*-values are shown for each.

Figure S2 Gene expression during proximal nephron formation in human nephrons and kidney organoids.

**a**. Dot plot of genes from an *in vivo* Pearson correlation test showing upregulation with human *HNF1B* (bottom), both *HNF1B* and *HNF4A* (middle), and just *HNF4A* (top). Clusters depict proximalizing lineages from the human nephron (left) and four kidney organoid datasets (right) using standard protocols. Green highlight denotes genes not upregulated in proximalizing organoid cells, red marks genes expressed in the distal nephron, and bolded genes are referenced in Figure S5. **b**. Venn diagram showing the intersection of gene lists from genes enriched in the adult proximal tubule (Kidney Cell Explorer; Ransick et al., 2019 *Dev Cell*), and genes significantly downregulated in the Hnf4a-mutant mouse (257 genes, *p*_adj_ < 0.01, log2FC ≥ 1.5; re-analyzed from Marable et al., 2020 *JASN*). Intersect categorizes Hnf4a-dependent genes enriched in development (124, right) and those enriched in development and adults (133, intersect). **c-d**. Heatmaps of human orthologs (generated using Ensembl BioMart) corresponding to the 133 mouse genes enriched in development and adult (c), and the 124 mouse genes enriched in development (d) from (b). Heatmaps indicate the percentage of cells expressing each gene in *HNF4A*^+^ clusters from human single-cell RNA-sequencing (Figure S1a) and kidney organoid nephron single-cell RNA-sequencing from multiple sources (Figure S1d). Scales at the bottom depict whether a gene is expressed at a higher percentage in human cells (blue) or in organoid nephron cells (red). Bolded genes are referenced in feature plots in (e) and (f). **e-f**. Feature plots displaying representative genes for each heatmap. Human insets highlight the transition from pretubular aggregate to later proximal nephron stages, while the organoid inset shows the proximalizing branch.

Figure S3 Immunofluorescent analyses of inhibitor-treated kidney organoids.

**a**. Whole-mount immunofluorescent stain of organoid nephrons on day 12 following 48 hr. culture in control conditions (untreated), LY294002-treated, DAPT-treated, and LY294002 + DAPT-treated. Insets highlight clusters of forming organoid nephrons, with split channels. Scale bars: 10 microns. **b**. Whole-mount immunofluorescent stains of days 13 and 14 control and LY294002-treated (from days 10-12, proximal-biased) organoids. Scale bars: 500 microns. **c**. Quantification of central and peripheral HNF4A^+^ and POU3F3^+^ organoid nephrons for control or proximal-biased conditions. The yellow line represents the organoid diameter, with the yellow circle marking the center of the organoid, and its diameter representing the radius of the whole organoid. Blue counts denote peripheral organoid nephrons, while red counts indicate central organoid nephrons. Organoids shown for this panel are the same that were profiled from Figure 2f. Scale bars: 200 microns. **d**. Percent contribution quantifications derived from (c). Bar graphs quantify structures out of the total number of central or peripheral structures as either HNF4A^+^ or POU3F3^+^. Data are averaged from *n* = 3 biological replicate kidney organoids.

Figure S4 Extended single-cell transcriptomics-driven analyses and comparison of kidney organoid nephrons with in vivo human nephrons.

**a**. Schematic of single-cell RNA-sequencing timepoints with UMAP reduction of kidney organoid cells, colored by samples. **b**. UMAP reduction of cell clusters in kidney organoids, colored by clusters. **c**. Dot plot with identifying genes per cluster for kidney organoid nephrogenic cells in Figure 3b. **d**. UMAP reduction of kidney organoid nephrogenic cells with podocytes removed, used for psuedotime trajectory analysis. **e**. Pseudotime trajectory of organoid nephron cells (with podocytes removed) from day 10 to day 18 proximal cells. **f**. Feature plots for representative genes marking different developmental stages from the pseudotime trajectory analysis. **g**. Single-cell RNA-sequencing UMAP reduction of human fetal nephron cells (Figure S1a) merged with kidney organoid nephrogenic cells (Figure 3b), colored by cluster. **h**. UMAP reduction from (g) colored by sample. **i**. Percent contribution bar graph quantifying the contribution of different original sample identities to various clusters of integrated nephrogenic cells from (g). **j**. Dot plot with representative genes for each cluster for human-organoid merged data in (g). **k**. Masking (red) of WT1^+^ organoid nephron segments from control and proximal-biased organoids on days 12 and 18 of differentiation. Scale bars: 200 microns. **l**. Quantification of the average size (µm^2^) of WT1^+^ organoid nephron segments from control and proximal-biased organoids on days 12 and 18 of differentiation, from (k). Data are quantified from *n* = 2 biological replicates each. SEM error bars are shown. Statistical significance is determined using a Mann-Whitney U test.

Figure S5 Single-cell transcriptomics-driven comparison of in vitro human proximalization with proximal-biased kidney organoid nephrons and proximal tubule-enhanced kidney organoids.

**a**. Single-cell RNA-sequencing of differentiation day 13+14 (27) proximal tubule-enhanced kidney organoids acquired from Vanslambrouck et al., 2022 *Nature Communications*. **b**. Dot plot with identifying genes for each cluster. **c**. Feature plots for select genes, showing a matching comparison to Figure S1c and S1f. **d**. Schematic of merging single-cell RNA-sequencing datasets: *in vivo* developing human nephrons from week 14 and 17 kidneys (Lindström et al., 2021 *Dev Cell*; Tran et al., 2019 *Dev Cell*) (Figure S1a), proximal-biased kidney organoids from differentiation days 10, 12, 14, and 18 (Figure 3a-b), and differentiation day 13+14 (27) proximal-tubule enhanced kidney organoids in (a). **e**. Diagram of differentiation and sampling timelines for proximal-biased kidney organoids and proximal tubule-enhanced kidney organoids. **f**. Violin plots of representative early proximal tubule genes (expressed in human clusters 21, 24, and 20, Figure S1a), split by dataset of origin. **g**. Violin plots of representative late proximal tubule genes (not expressed in human cluster 21, expressed in clusters 24 and 20, Figure S1a), split by dataset of origin.

Figure S6 Dynamic imaging, functional profiling, and reproducibility of kidney organoid proximal biasing.

**a**. Comparative live imaging of HNF4A-YFP reporter (Vanslambrouck et al., 2019 *JASN*) kidney organoids for control vs. proximal-biased conditions, spanning differentiation days 10 to 18. Scale bars: 200 microns. **b**. Live imaging of HNF4A-YFP kidney organoids on differentiation day 18 with TRITC-albumin uptake. Quantification of individual uptake events displayed in lower left, and insets detail HNF4A-YFP and TRITC-albumin distinction. Scale bars: whole organoid-500 microns, insets-100 microns. **c**. Quantitative data from organoid nephron single-cell RNA-sequencing (Figure 3a-b) showing cell counts in HNF4A^+^ proximal clusters, categorized by control or proximal-biased conditions, expressing select genes. **d**. Whole-mount immunofluorescence comparing control and proximal-biased day 18 kidney organoids from an alternate iPSC line. Scale bars: 10 microns.

## REFERENCES

1. Zhuo, J. L. & Li, X. C. Proximal nephron. Compr. Physiol. 3, 1079–123 (2013).

2. Chevalier, R. L. The proximal tubule is the primary target of injury and progression of kidney disease: role of the glomerulotubular junction. Am. J. Physiol. Renal Physiol. 311, F145–61 (2016).

3. Nakhoul, N. & Batuman, V. Role of proximal tubules in the pathogenesis of kidney disease. Contrib. Nephrol. 169, 37–50 (2011).

4. Takasato, M., Er, P. X., Chiu, H. S. & Little, M. H. Generation of kidney organoids from human pluripotent stem cells. Nat. Protoc. 11, 1681–1692 (2016).

5. Morizane, R. & Bonventre, J. V. Generation of nephron progenitor cells and kidney organoids from human pluripotent stem cells. Nat. Protoc. 12, 195–207 (2017).

6. Combes, A. N., Zappia, L., Er, P. X., Oshlack, A. & Little, M. H. Single-cell analysis reveals congruence between kidney organoids and human fetal kidney. Genome Med. 11, 3 (2019).

7. Subramanian, A. et al. Single cell census of human kidney organoids shows reproducibility and diminished off-target cells after transplantation. Nat. Commun. 10, 5462 (2019).

8. Tran, T. et al. In Vivo Developmental Trajectories of Human Podocyte Inform In Vitro Differentiation of Pluripotent Stem Cell-Derived Podocytes. Dev. Cell 50, 102–116.e6 (2019).

9. Wu, H. et al. Comparative Analysis and Refinement of Human PSC-Derived Kidney Organoid Differentiation with Single-Cell Transcriptomics. Cell Stem Cell 23, 869–881.e8 (2018).

10. Vanslambrouck, J. M. et al. Enhanced metanephric specification to functional proximal tubule enables toxicity screening and infectious disease modelling in kidney organoids. Nat. Commun. 13, 5943 (2022).

11. Yoshimura, Y. et al. A single-cell multiomic analysis of kidney organoid differentiation. Proc. Natl. Acad. Sci. U. S. A. 120, e2219699120 (2023).

12. Morizane, R. et al. Nephron organoids derived from human pluripotent stem cells model kidney development and injury. Nat. Biotechnol. 33, 1193–200 (2015).

13. Gupta, N. et al. Modeling injury and repair in kidney organoids reveals that homologous recombination governs tubular intrinsic repair. Sci. Transl. Med. 14, eabj4772 (2022).

14. Digby, J. L. M., Vanichapol, T., Przepiorski, A., Davidson, A. J. & Sander, V. Evaluation of cisplatin-induced injury in human kidney organoids. Am. J. Physiol. Renal Physiol. 318, F971–F978 (2020).

15. Yoshimura, Y., Muto, Y., Omachi, K., Miner, J. H. & Humphreys, B. D. Elucidating the Proximal Tubule HNF4A Gene Regulatory Network in Human Kidney Organoids. J. Am. Soc. Nephrol. JASN (2023) doi:10.1681/ASN.0000000000000197.

16. Marable, S. S., Chung, E. & Park, J.-S. Hnf4a Is Required for the Development of Cdh6-Expressing Progenitors into Proximal Tubules in the Mouse Kidney. J. Am. Soc. Nephrol. JASN 31, 2543–2558 (2020).

17. Marable, S. S., Chung, E., Adam, M., Potter, S. S. & Park, J.-S. Hnf4a deletion in the mouse kidney phenocopies Fanconi renotubular syndrome. JCI Insight 3, (2018).

18. Lindström, N. O. et al. Spatial transcriptional mapping of the human nephrogenic program. Dev. Cell 56, 2381–2398.e6 (2021).

19. Schnell, J., Achieng, M. & Lindström, N. O. Principles of human and mouse nephron development. Nat. Rev. Nephrol. (2022) doi:10.1038/s41581-022-00598-5.

20. Georgas, K. et al. Analysis of early nephron patterning reveals a role for distal RV proliferation in fusion to the ureteric tip via a cap mesenchyme-derived connecting segment. Dev. Biol. 332, 273–86 (2009).

21. Lindström, N. O. et al. Progressive Recruitment of Mesenchymal Progenitors Reveals a Time-Dependent Process of Cell Fate Acquisition in Mouse and Human Nephrogenesis. Dev. Cell 45, 651–660.e4 (2018).

22. Chung, E., Deacon, P. & Park, J.-S. Notch is required for the formation of all nephron segments and primes nephron progenitors for differentiation. Dev. Camb. Engl. 144, 4530–4539 (2017).

23. Heliot, C. et al. HNF1B controls proximal-intermediate nephron segment identity in vertebrates by regulating Notch signalling components and Irx1/2. Dev. Camb. Engl. 140, 873–85 (2013).

24. Massa, F. et al. Hepatocyte nuclear factor 1β controls nephron tubular development. Dev. Camb. Engl. 140, 886–96 (2013).

25. Lindström, N. O. et al. Integrated β-catenin, BMP, PTEN, and Notch signalling patterns the nephron. eLife 3, e04000 (2015).

26. Ng-Blichfeldt, J.-P., Stewart, B. J., Clatworthy, M. R., Williams, J. M. & Röper, K. Identification of a core transcriptional program driving the human renal mesenchymal-to-epithelial transition. Dev. Cell (2024) doi:10.1016/j.devcel.2024.01.011.

27. Ransick, A. et al. Single-Cell Profiling Reveals Sex, Lineage, and Regional Diversity in the Mouse Kidney. Dev. Cell 51, 399–413.e7 (2019).

28. Love, M. I., Huber, W. & Anders, S. Moderated estimation of fold change and dispersion for RNA-seq data with DESeq2. Genome Biol. 15, 550 (2014).

29. Lindström, N. O., Carragher, N. O. & Hohenstein, P. The PI3K pathway balances self-renewal and differentiation of nephron progenitor cells through β-catenin signaling. Stem Cell Rep. 4, 551–60 (2015).

30. Vlahos, C. J., Matter, W. F., Hui, K. Y. & Brown, R. F. A specific inhibitor of phosphatidylinositol 3-kinase, 2-(4-morpholinyl)-8-phenyl-4H-1-benzopyran-4-one (LY294002). J. Biol. Chem. 269, 5241–8 (1994).

31. Wang, J. et al. PI3K-AKT pathway mediates growth and survival signals during development of fetal mouse lung. Tissue Cell 37, 25–35 (2005).

32. Duvall, K. et al. Revisiting the role of notch in nephron segmentation confirms a role for proximal fate selection during mouse and human nephrogenesis. Dev. Camb. Engl. (2022) doi:10.1242/dev.200446.

33. Cheng, H.-T. et al. Notch2, but not Notch1, is required for proximal fate acquisition in the mammalian nephron. Dev. Camb. Engl. 134, 801–11 (2007).

34. Dovey, H. F. et al. Functional gamma-secretase inhibitors reduce beta-amyloid peptide levels in brain. J. Neurochem. 76, 173–81 (2001).

35. Cheng, H.-T. et al. Gamma-secretase activity is dispensable for mesenchyme-to-epithelium transition but required for podocyte and proximal tubule formation in developing mouse kidney. Dev. Camb. Engl. 130, 5031–42 (2003).

36. Folkes, A. J. et al. The identification of 2-(1H-indazol-4-yl)-6-(4-methanesulfonyl-piperazin-1-ylmethyl)-4-morpholin-4-yl-thieno[3,2-d]pyrimidine (GDC-0941) as a potent, selective, orally bioavailable inhibitor of class I PI3 kinase for the treatment of cancer. J. Med. Chem. 51, 5522–32 (2008).

37. Lindström, N. O. et al. Conserved and Divergent Features of Mesenchymal Progenitor Cell Types within the Cortical Nephrogenic Niche of the Human and Mouse Kidney. J. Am. Soc. Nephrol. JASN 29, 806–824 (2018).

38. Haghverdi, L., Lun, A. T. L., Morgan, M. D. & Marioni, J. C. Batch effects in single-cell RNA-sequencing data are corrected by matching mutual nearest neighbors. Nat. Biotechnol. 36, 421–427 (2018).

39. Ren, Q. et al. Distinct functions of megalin and cubilin receptors in recovery of normal and nephrotic levels of filtered albumin. Am. J. Physiol. Renal Physiol. 318, F1284–F1294 (2020).

40. Bajaj, P. et al. Human Pluripotent Stem Cell-Derived Kidney Model for Nephrotoxicity Studies. Drug Metab. Dispos. Biol. Fate Chem. 46, 1703–1711 (2018).

41. Schuh, C. D. et al. Combined Structural and Functional Imaging of the Kidney Reveals Major Axial Differences in Proximal Tubule Endocytosis. J. Am. Soc. Nephrol. JASN 29, 2696–2712 (2018).

42. Molitoris, B. A., Sandoval, R. M., Yadav, S. P. S. & Wagner, M. C. Albumin uptake and processing by the proximal tubule: physiological, pathological, and therapeutic implications. Physiol. Rev. 102, 1625–1667 (2022).

43. Vanslambrouck, J. M. et al. A Toolbox to Characterize Human Induced Pluripotent Stem Cell-Derived Kidney Cell Types and Organoids. J. Am. Soc. Nephrol. JASN 30, 1811–1823 (2019).

44. Pabla, N., Murphy, R. F., Liu, K. & Dong, Z. The copper transporter Ctr1 contributes to cisplatin uptake by renal tubular cells during cisplatin nephrotoxicity. Am. J. Physiol. Renal Physiol. 296, F505–11 (2009).

45. Miller, R. P., Tadagavadi, R. K., Ramesh, G. & Reeves, W. B. Mechanisms of Cisplatin nephrotoxicity. Toxins 2, 2490–518 (2010).

46. Motwani, S. S., Kaur, S. S. & Kitchlu, A. Cisplatin Nephrotoxicity: Novel Insights Into Mechanisms and Preventative Strategies. Semin. Nephrol. 42, 151341 (2022).

47. Tang, C., Livingston, M. J., Safirstein, R. & Dong, Z. Cisplatin nephrotoxicity: new insights and therapeutic implications. Nat. Rev. Nephrol. 19, 53–72 (2023).

48. Yamashita, N. et al. Cumulative DNA damage by repeated low-dose cisplatin injection promotes the transition of acute to chronic kidney injury in mice. Sci. Rep. 11, 20920 (2021).

49. Huang, G., Zhang, Q., Xu, C., Chen, L. & Zhang, H. Mechanism of kidney injury induced by cisplatin. Toxicol. Res. 11, 385–390 (2022).

50. Singh, G. A possible cellular mechanism of cisplatin-induced nephrotoxicity. Toxicology 58, 71–80 (1989).

51. Sánchez-González, P. D., López-Hernández, F. J., López-Novoa, J. M. & Morales, A. I. An integrative view of the pathophysiological events leading to cisplatin nephrotoxicity. Crit. Rev. Toxicol. 41, 803–21 (2011).

52. Choie, D. D., Longnecker, D. S. & del Campo, A. A. Acute and chronic cisplatin nephropathy in rats. Lab. Investig. J. Tech. Methods Pathol. 44, 397–402 (1981).

53. Cullen, K. J., Yang, Z., Schumaker, L. & Guo, Z. Mitochondria as a critical target of the chemotheraputic agent cisplatin in head and neck cancer. J. Bioenerg. Biomembr. 39, 43– 50 (2007).

54. Mandic, A., Hansson, J., Linder, S. & Shoshan, M. C. Cisplatin induces endoplasmic reticulum stress and nucleus-independent apoptotic signaling. J. Biol. Chem. 278, 9100–6 (2003).

55. Gerhardt, L. M. S., Liu, J., Koppitch, K., Cippà, P. E. & McMahon, A. P. Single-nuclear transcriptomics reveals diversity of proximal tubule cell states in a dynamic response to acute kidney injury. Proc. Natl. Acad. Sci. U. S. A. 118, (2021).

56. Gerhardt, L. M. S. et al. Lineage Tracing and Single-Nucleus Multiomics Reveal Novel Features of Adaptive and Maladaptive Repair after Acute Kidney Injury. J. Am. Soc. Nephrol. JASN (2023) doi:10.1681/ASN.0000000000000057.

57. Ledru, N. et al. Predicting proximal tubule failed repair drivers through regularized regression analysis of single cell multiomic sequencing. Nat. Commun. 15, 1291 (2024).

58. Aggarwal, S. et al. SOX9 switch links regeneration to fibrosis at the single-cell level in mammalian kidneys. Science 383, eadd6371 (2024).

59. Kumar, S. et al. Sox9 Activation Highlights a Cellular Pathway of Renal Repair in the Acutely Injured Mammalian Kidney. Cell Rep. 12, 1325–1338 (2015).

60. Kang, H. M. et al. Sox9-Positive Progenitor Cells Play a Key Role in Renal Tubule Epithelial Regeneration in Mice. Cell Rep. 14, 861–871 (2016).

61. Zhang, K. et al. In vivo two-photon microscopy reveals the contribution of Sox9+ cell to kidney regeneration in a mouse model with extracellular vesicle treatment. J. Biol. Chem. 295, 12203–12213 (2020).

62. Chen, L. & Al-Awqati, Q. Segmental expression of Notch and Hairy genes in nephrogenesis. Am. J. Physiol. Renal Physiol. 288, F939–52 (2005).

63. Liu, Z. et al. The extracellular domain of Notch2 increases its cell-surface abundance and ligand responsiveness during kidney development. Dev. Cell 25, 585–98 (2013).

64. Bonegio, R. G. B., Beck, L. H., Kahlon, R. K., Lu, W. & Salant, D. J. The fate of Notch-deficient nephrogenic progenitor cells during metanephric kidney development. Kidney Int. 79, 1099–112 (2011).

65. Bohn, S. et al. Distinct molecular and morphogenetic properties of mutations in the human HNF1beta gene that lead to defective kidney development. J. Am. Soc. Nephrol. JASN 14, 2033–41 (2003).

66. Nakayama, M. et al. HNF1B alterations associated with congenital anomalies of the kidney and urinary tract. Pediatr. Nephrol. Berl. Ger. 25, 1073–9 (2010).

67. Naylor, R. W., Przepiorski, A., Ren, Q., Yu, J. & Davidson, A. J. HNF1β is essential for nephron segmentation during nephrogenesis. J. Am. Soc. Nephrol. JASN 24, 77–87 (2013).

68. Clissold, R. L., Hamilton, A. J., Hattersley, A. T., Ellard, S. & Bingham, C. HNF1B-associated renal and extra-renal disease-an expanding clinical spectrum. Nat. Rev. Nephrol. 11, 102– 12 (2015).

69. Przepiorski, A. et al. A Simple Bioreactor-Based Method to Generate Kidney Organoids from Pluripotent Stem Cells. Stem Cell Rep. 11, 470–484 (2018).

70. Chen, L. et al. A reinforcing HNF4-SMAD4 feed-forward module stabilizes enterocyte identity. Nat. Genet. 51, 777–785 (2019).

71. Chen, L., et al. HNF4 factors control chromatin accessibility and are redundantly required for maturation of the fetal intestine. Dev. Camb. Engl. 146, dev179432 (2019).

72. Guo, Q. et al. A β-catenin-driven switch in tcf/lef transcription factor binding to dna target sites promotes commitment of mammalian nephron progenitor cells. eLife 10, 1–25 (2021).

73. Park, J.-S. et al. Six2 and Wnt regulate self-renewal and commitment of nephron progenitors through shared gene regulatory networks. Dev. Cell 23, 637–51 (2012).

74. Lindström, N. O. et al. Conserved and Divergent Molecular and Anatomic Features of Human and Mouse Nephron Patterning. J. Am. Soc. Nephrol. JASN 29, 825–840 (2018).

75. Blaine, J., Chonchol, M. & Levi, M. Renal control of calcium, phosphate, and magnesium homeostasis. Clin. J. Am. Soc. Nephrol. CJASN 10, 1257–72 (2015).

76. Klootwijk, E. D., et al. Renal Fanconi syndrome: taking a proximal look at the nephron. Nephrol. Dial. Transplant. Off. Publ. Eur. Dial. Transpl. Assoc.-Eur. Ren. Assoc. 30, 1456– 60 (2015).

77. Baum, M. & Quigley, R. Proximal tubule water transport-lessons from aquaporin knockout mice. Am. J. Physiol. Renal Physiol. 289, F1193–4 (2005).

78. Wilmes, A. et al. Mechanism of cisplatin proximal tubule toxicity revealed by integrating transcriptomics, proteomics, metabolomics and biokinetics. Toxicol. Vitro Int. J. Publ. Assoc. BIBRA 30, 117–27 (2015).

79. Nieskens, T. T. G. et al. Expression of Organic Anion Transporter 1 or 3 in Human Kidney Proximal Tubule Cells Reduces Cisplatin Sensitivity. Drug Metab. Dispos. Biol. Fate Chem. 46, 592–599 (2018).

80. Cippà, P. E. & McMahon, A. P. Proximal tubule responses to injury: interrogation by single-cell transcriptomics. Curr. Opin. Nephrol. Hypertens. (2023) doi:10.1097/MNH.0000000000000893.

81. Banan Sadeghian, R., et al. Cells sorted off hiPSC-derived kidney organoids coupled with immortalized cells reliably model the proximal tubule. *Commun*. Biol. 6, 483 (2023).

82. Yousef Yengej, F. A., et al. Tubuloid culture enables long-term expansion of functional human kidney tubule epithelium from iPSC-derived organoids. Proc. Natl. Acad. Sci. U. S. A. 120, e2216836120 (2023).

83. Aceves, J. O. et al. 3D proximal tubule-on-chip model derived from kidney organoids with improved drug uptake. Sci. Rep. 12, 14997 (2022).

84. Carracedo, M. et al. 3D vascularised proximal tubules-on-a-multiplexed chip model for enhanced cell phenotypes. Lab. Chip (2023) doi:10.1039/d2lc00723a.

85. Howden, S. E. et al. A Cas9 Variant for Efficient Generation of Indel-Free Knockin or Gene-Corrected Human Pluripotent Stem Cells. Stem Cell Rep. 7, 508–517 (2016).

86. Takasato, M. & Little, M. H. A strategy for generating kidney organoids: Recapitulating the development in human pluripotent stem cells. Dev. Biol. 420, 210–220 (2016).

87. Howden, S. E. & Little, M. H. Generating Kidney Organoids from Human Pluripotent Stem Cells Using Defined Conditions. Methods Mol. Biol. Clifton NJ 2155, 183–192 (2020).

88. Stuart, T. et al. Comprehensive Integration of Single-Cell Data. Cell (2019) doi:10.1016/j.cell.2019.05.031.

89. Samuel Marsh, Maëlle Salmon & Paul Hoffman. samuel-marsh/scCustomize: Version 2.1.2. [object Object] 10.5281/ZENODO.5706430 (2024).

90. Luecken, M. D. et al. Benchmarking atlas-level data integration in single-cell genomics. Nat. Methods 19, 41–50 (2022).

91. Trapnell, C. et al. The dynamics and regulators of cell fate decisions are revealed by pseudotemporal ordering of single cells. Nat. Biotechnol. 32, 381–386 (2014).

92. Qiu, X. et al. Reversed graph embedding resolves complex single-cell trajectories. Nat. Methods 14, 979–982 (2017).

93. Qiu, X. et al. Single-cell mRNA quantification and differential analysis with Census. Nat. Methods 14, 309–315 (2017).

94. Cao, J. et al. The single-cell transcriptional landscape of mammalian organogenesis. Nature 566, 496–502 (2019).

95. Dobin, A. et al. STAR: ultrafast universal RNA-seq aligner. Bioinforma. Oxf. Engl. 29, 15–21 (2013).

96. Schindelin, J., et al. Fiji: an open-source platform for biological-image analysis. Nat. Methods 9, 676–82 (2012).

